# Contractile perinuclear actomyosin network promotes peripheral and polar chromosome interaction with the mitotic spindle

**DOI:** 10.1101/2025.09.19.677380

**Authors:** Nooshin Sheidaei, John K. Eykelenboom, Zuojun Yue, Graeme Ball, Alexander J.R. Booth, Tomoyuki U. Tanaka

## Abstract

Chromosomes must efficiently and properly interact with the mitotic spindle during prometaphase for correct segregation in anaphase. Chromosomes at the nuclear periphery or behind the spindle poles interact less efficiently with the mitotic spindle, increasing the risk of missegregation. The mechanisms that mitigate such risks in unperturbed cells are unknown. An actomyosin network (PANEM) forms around the nucleus during prophase. While the myosin-II-dependent PANEM contraction immediately after nuclear envelope breakdown (NEBD) facilitates chromosome interaction with the mitotic spindle, the mechanism by which it does so remains unclear. Here, using human cell lines, we show that immediately after NEBD, PANEM contraction directly pushes chromosomes at the nuclear periphery or behind spindle poles toward the center of cells. Detailed tracking of kinetochore movements following light-induced activation of a myosin-II inhibitor reveals that this inward movement of chromosomes facilitates kinetochores’ initial interaction with spindle microtubules. It also promotes the onset of kinetochores’ congression towards the spindle mid-plane, but not congression itself once it starts. Thus, PANEM contraction ensures high-fidelity chromosome segregation by relocating chromosomes from unfavorable locations. Since some chromosomally unstable cancer cells fail to establish PANEM during early mitosis, the absence of PANEM may contribute to numerical chromosomal instability in these cells.

**Impact statement:** An actomyocin network (PANEM) forms around the nucleus in prophase, and its contraction repositions peripheral and polar chromosomes to facilitate their interaction with the mitotic spindle, ensuring their correct segregation.

## Introduction

For high-fidelity chromosome segregation in human cells, all chromosomes must correctly interact with microtubules (MTs) of the mitotic spindle during the early stage of mitosis (prometaphase), i.e. shortly after nuclear envelope breakdown (NEBD). To facilitate this, the bipolar mitotic spindle is established when two spindle poles separate from each other around the time of NEBD ^1^, while the kinetochore provides the major MT interaction site on a chromosome ^2,3,4^. The kinetochore interacts with MTs emanating from spindle poles in a stepwise manner ^5,6^: it initially interacts with the lateral side of a MT and then travels along it towards a spindle pole, before it attaches to the plus end of the same MT after this MT shrinks ^7–10,11^. The other sister kinetochore then interacts with the side of another MT (either another kinetochore-attached MT or a pole-to-pole MT) and moves along it towards the middle of the mitotic spindle (spindle mid-plane or metaphase plate) in a process called chromosome congression ^12,13^. Finally, biorientation is established when one of the sister kinetochores interacts with MTs only from one spindle pole and the other sister kinetochore with MTs only from the opposite pole. ^6,14^, leading to the oscillatory motions of chromosomes around the spindle mid-plane ^15–17^.

For efficient and correct kinetochore-MT interaction, the locations of chromosomes in the nucleus (when NEBD occurs) or relative to spindle poles (after NEBD occurs) are crucial. For example, in cancerous cells, or non-transformed cells treated with a low-level Mps1 inhibitor, chromosomes at the nuclear periphery or behind spindle poles (polar regions) early in mitosis are more likely to missegregate during anaphase than internally located chromosomes^18,19^. In other words, these polar/peripheral chromosome locations in early mitosis are unfavorable for correct segregation later in mitosis. It is thought that (1) chromosomes at the nuclear periphery are located further away from spindle poles and therefore require a longer time to interact with spindle MTs extending from spindle poles, and (2) chromosomes at polar regions require a longer time to start congression to the spindle mid-plane ^18,19^. Nevertheless, the missegregation rate of these chromosomes is still relatively low in unperturbed normal cells, meaning there might be mechanisms mitigating the risks of missegregation for chromosomes found at these locations. However, such mechanisms have not yet been identified.

Recently, potentially relevant to such mechanisms, a LINC complex-dependent actomyosin network that is rapidly formed on the cytoplasmic side of the nuclear envelope (NE) during prophase was reported ^20,21^ (Figure 1A). In the current study, we call this actomyosin network PANEM (**P**erinuclear **A**ctomyosin **N**etwork in **E**arly **M**itosis). The PANEM is formed in U2OS and RPE1 cells, but not in HeLa cells ^20,21^. In prophase, the PANEM facilitates the separation of spindle poles to establish a bipolar spindle ^21^. Following NEBD, the PANEM remains on the NE remnants for 10-15 min and shows contraction promoted by myosin II ^20^ (Figure 1A). Concomitantly, chromosome scattering volume (CSV), quantified by three dimensional convex hull fitting ^20^, also becomes smaller (Figure 1B). The reduction in CSV after NEBD also occurred after MT disruption with nocodazole treatment, suggesting that it was not dependent on chromosome interaction with spindle MTs ^20^. It was concluded that the myosin II-dependent PANEM contraction promoted CSV reduction. Furthermore, in situations where PANEM contraction was inhibited, (1) chromosomes often remained in polar regions, (2) anaphase onset was frequently delayed, and (3) chromosome missegregation often occurred ^20^ (Figure 1 – figure supplement 1). The PANEM contraction seemed to facilitate correct chromosome interaction with spindle MTs. However, it has been unclear how PANEM contraction promotes the stepwise development of kinetochore interaction with spindle MTs mentioned above or which chromosomes specifically benefit from the PANEM contraction in this process.

**Figure 1:**
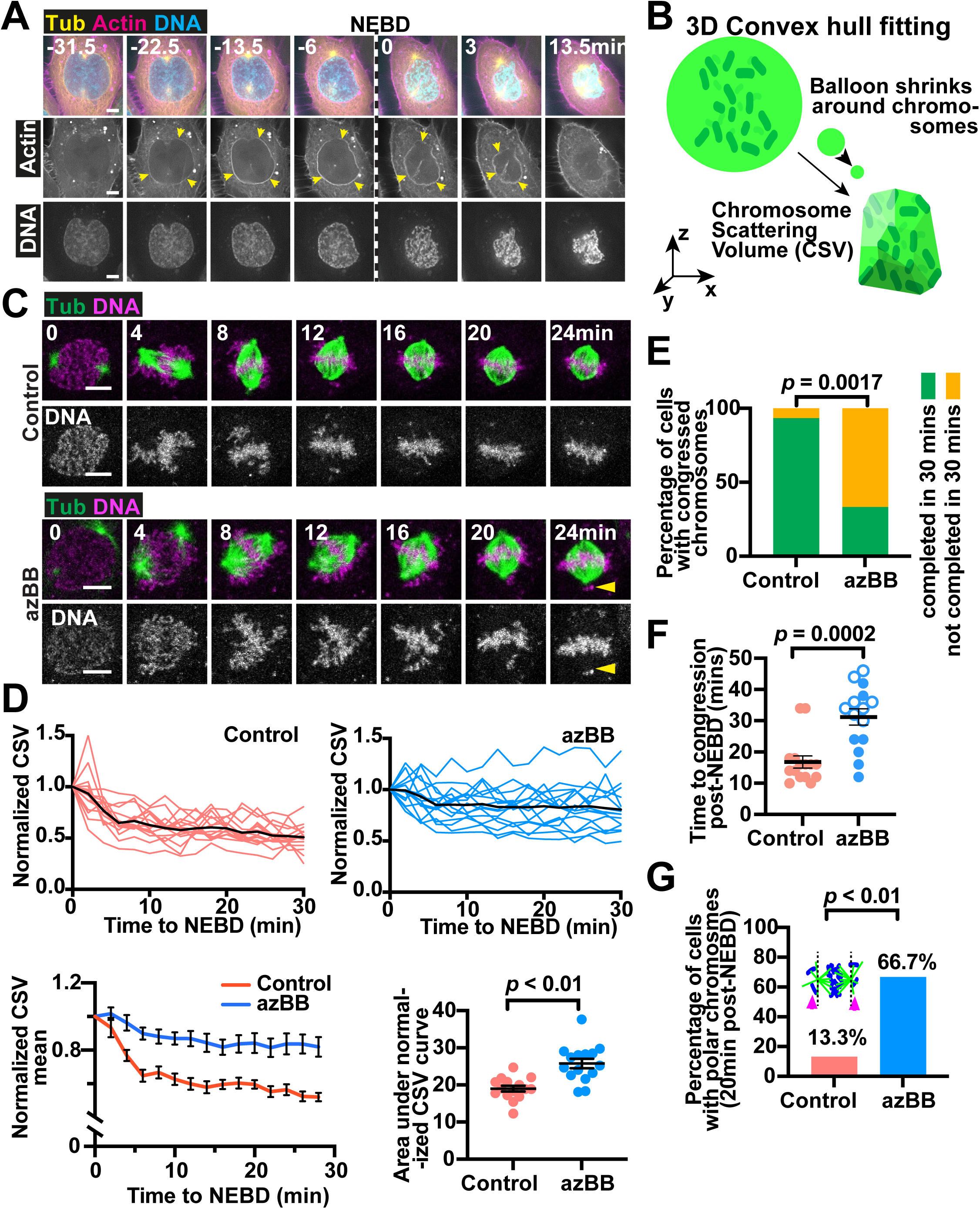
The outcome of selective and rapid suppression of perinuclear actomyosin network (PANEM) contraction. **(A)** Time-lapse images show a representative cell passing through the early stages of mitosis (prophase and prometaphase). A stable cell line expressing mCherry-LifeAct and GFP-⍰Tubulin, with chromosomes visualized by SYTO deep-red was imaged every 1.5 minutes. The timing of nuclear envelope breakdown (NEBD) is indicated by the dotted line. The PANEM is indicated by yellow arrowheads. Scale bar is 5µm. **(B)** Diagram shows how the chromosome scattering volume (CSV) was defined. The convex-hull was fitted in three dimensions (3D) to represent chromosome distribution. A hypothetical ‘balloon’ (green) shrinks around chromosomes (dark green) in 3D to create a convex hull or a minimal polyhedron wrapping chromosomes. The volume of the 3D convex hull indicates how widely chromosomes are scattered in space. **(C)** Time-lapse images show representative cells passing through mitosis (prometaphase to metaphase). A stable cell line expressing GFP-⍰Tubulin, with chromosomes visualized by SiRDNA, was imaged every 4 minutes following treatment with or without azBB followed by nuclear irradiation using a 2-photon 860nm laser. NEBD occurred at 0 minute and was detected by diffusion of free GFP-⍰Tubulin into the nuclear space. Chromosomes that did not congress and lie behind the spindle poles are indicated by yellow arrowheads. Scale bar is 10µm. Time-lapse images, exemplified here, were used in the analyses in D–G. **(D)** Graphs in the upper panels show normalized chromosome scattering volume (CSV; see B) before and after NEBD (0 min) for cells treated with or without azBB. The data is normalized to the volume at 0 minutes (immediately after NEBD). Each red or blue line represents the measurements from an individual cell while the heavy black lines represent the mean measurement across the time points. In the lower left panel the mean of normalized values obtained above are presented here again with standard error of mean (SEM) shown for each time point for each condition. In the lower right panel, the graph plot the areas under the curves for normalized CSV, measured in individual cells. The bars represent the mean and SEM. The *p* value was obtained using t-test. The data without normalization is shown in Figure 1 – figure supplement 3. **(E)** Graph shows the percentage of cells, treated either with or without azBB, whose chromosomes were congressed before 30min (green) or not (orange). The number of cells for each group was 15. The *p* value was obtained using Fisher’s exact test. **(F)** Graph shows the time (from NEBD) taken for completion of congression in individual cells treated either with or without azBB. Open circle data points represent cells that had not completed congression at the end of the time lapse sequence. The bars represent the mean and SEM. The *p* value obtained using an unpaired Mann-Whitney test. **(G)** Graph shows the percentage of cells which exhibited chromosomes behind the spindle poles 20 minutes after NEBD. The cartoon shows microtubules (green), boundaries for polar regions (the black dotted lines), chromosomes (blue), and polar chromosomes (pink arrowheads). The number of cells analyzed for each group was 15. The *p* value was obtained using Fisher’s exact test.

In this work, we address which steps of kinetochore-MT interaction are facilitated by the PANEM contraction and which kinetochores are helped by the PANEM contraction to establish correct MT interaction. We find that soon after NEBD, the PANEM contraction assists kinetochores at the nuclear periphery in initiating interaction with spindle MTs. Moreover, the PANEM contraction helps the kinetochores in polar regions (i.e. behind spindle poles) to start congression towards the middle of the mitotic spindle. The PANEM contraction has such effects because it pushes chromosomes inward, thus repositioning them to facilitate their correct interaction with spindle MTs. Therefore, PANEM contraction repositions chromosomes from unfavorable locations to establish their correct interaction with spindle MTs, thus mitigating the risks of their missegregation later in mitosis. In this study, we also broaden our analysis of PANEM to multiple human cell lines. While PANEM was also found to be required for reducing CSV in a human non-cancerous cell line, several cancer cells with high levels of aneuploidy were found to lack PANEM in early mitosis. This suggests a link between PANEM and numerical chromosomal instability (N-CIN), which is associated with cancer progression.

## Results

### Developing a method for selective and prompt suppression of the PANEM contraction soon after NEBD

We aimed to study how PANEM contraction facilitates chromosome interaction with spindle MTs, including their initial interaction. Our previous data and other studies suggest the initial chromosome interaction with spindle MTs occurs within 8 min of NEBD ^5,20,22–25^. Therefore, we aimed to suppress PANEM contraction within this time range and study the outcome. While the PANEM contour becomes ambiguous about 10 min after NEBD, the CSV reduction reflects the extent of PANEM contraction for a longer time ^20^ (see Introduction). To suppress the PANEM contraction and CSV reduction, we previously used para-nitro-blebbistatin (pnBB), an inhibitor of myosin-II ^20^. However, this suppression was only apparent after 8 min following NEBD. We suspected the delayed effect was due to the partial inhibition of the myosin II activity by pnBB.

In the current study, to suppress the myosin II activity more efficiently, we used azido-blebbistatin (azBB). When excited with two-photon infrared light, azBB covalently binds the heavy chain of myosin II and inhibits its ATPase activity ^26,27^. We used a low concentration of azBB to ensure that myosin II was inhibited only at the region exposed to infrared light in U2OS cells. To inhibit myosin II on the PANEM, only the nucleus was irradiated by the infrared light during prophase, avoiding the cell cortex where a high amount of myosin II localizes ^28^. In the control, the nucleus was also irradiated by infrared light, but without azBB treatment. We confirmed that cell rounding during mitosis, which requires myosin II at the cell cortex ^28^, occurred similarly between the azBB treatment and the control (Figure 1 – figure supplement 2). This suggests that, as intended, myosin II of the cell cortex remained functional.

To enrich prophase cells, we arrested U2OS *cdk1-as* cells (with GFP-⍰Tubulin) at the end of G2 phase by treating them with an ATP analogue 1NM-PP1 ^20,21^. After 1NM-PP1 washout, these cells entered mitosis synchronously, and chromosomes were visualized with SiRDNA. The nuclei of prophase cells (those showing partial chromosome condensation) were exposed to infrared light in the presence and absence of azBB (Figure 1C), and the change of CSV was measured over time (Figure 1D, Figure 1 – figure supplement 3). NEBD was identified by the influx of GFP-⍰Tubulin signals (not incorporated into MTs) from the cytoplasm into the nucleus. In the control cells (without azBB), CSV reduction occurred rapidly until 6 min after NEBD and more slowly for the next 10 min or so (Figure 1D) – these results are consistent with the previous report ^20^. With azBB, the CSV reduction was weakened, compared with control cells – the difference was significant from 4 min after NEBD and onwards (Figure 1D). Treatment of cells with nocodazole (with and without azBB) before irradiation did not affect the outcomes, suggesting that the effect of azBB was independent of MTs (Figure 1 – figure supplement 4). Moreover, we also measured the volume inside the PANEM with and without azBB – the reduction of this volume was also weakened with azBB treatment within 8 min after NEBD (Figure 1 – figure supplement 5).

Furthermore, we previously showed that the suppression of the PANEM contraction with pnBB led to (1) a delay in chromosome congression and anaphase onset, (2) an increase in chromosomes at the polar region (behind spindle poles), leading to chromosome missegregation ^20^. These defects were also observed after the PANEM contraction was suppressed with azBB treatment (Figure 1E-G). It was previously shown that the PANEM has a role in spindle pole positioning during mitosis ^21^. Therefore, the problems we observed with azBB might have resulted from spindle assembly defects. However, the distribution of spindle length in control and azBB-treated cells was similar (Figure 1 – figure supplement 6), indicating that the spindle assembly was largely normal after the PANEM contraction was suppressed with azBB.

In summary, both the PANEM contraction and consequent CSV reduction were dependent on the activity of myosin II, and both occurred shortly (within 8 min) after NEBD. Both events can be suppressed by treatment with the myosin II inhibitor azBB within this time range. Therefore, we decided to use azBB to rapidly suppress the PANEM contraction following NEBD and study how this affects chromosome interactions with spindle MTs (see below).

### Kinetochore-MT interaction is developed through four distinct phases during prometaphase

The kinetochore provides the main MT attachment site on a chromosome. To investigate the effect of PANEM contraction on kinetochore-MT interaction, we first sought to distinguish the sequential phases through which this interaction is developed and established. For this, U2OS cells were synchronized as in the previous section, and we imaged the kinetochore (CENP-B-mCherry) and MTs (GFP-⍰Tubulin) every 30 s. We were able to track the positions of individual kinetochores over time if they were not too crowded (Figure 2 – figure supplement 1). We then tracked the position of each of these kinetochores, over time, relative to the following three reference points: (1) the midpoint between spindle poles, (2) the mid-plane between spindle poles, and (3) a spindle pole (the one towards which the kinetochore moved after NEBD) (Figure 2A).

**Figure 2:**
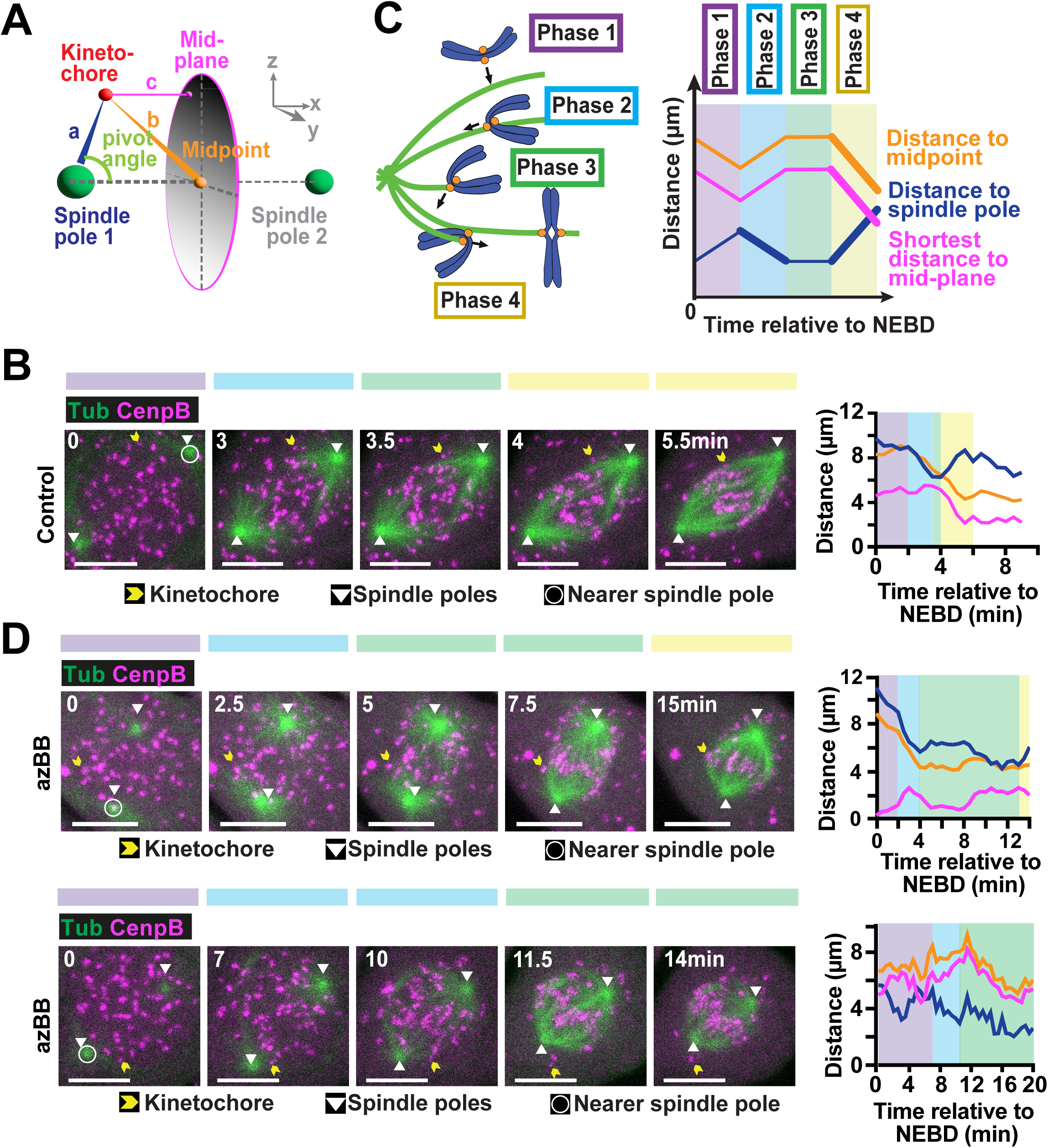
Tracking kinetochores during early phases of mitosis reveals four major phases of motion leading to congression to the spindle mid-plane. **(A)** To assess kinetochore motions during early mitosis (prometaphase), distances were calculated between the kinetochore and the nearest spindle pole, the spindle mid-point and the metaphase plate (mid-plane). To assess the location of the kinetochore relative to the spindle poles (polar or non-polar), the pivot angle was calculated. Designated color codes for each distance are indicated in the diagram. **(B)** Time-lapse images show a representative control cell passing through the early stages of mitosis (prometaphase). A stable cell line expressing CENPB-mCherry and GFP-⍰Tubulin was imaged every 30 seconds. The spindle poles are indicated by white arrowheads. The tracked kinetochore is indicated in each frame by a yellow arrowhead. The nearest spindle pole is indicated by a white circle in the first image of the sequence. Scale bar is 10µm. The graph on the right-hand side indicates the distance of the indicated kinetochore to the nearest spindle pole (dark blue line), the spindle midpoint (orange line) and the spindle mid-plane (magenta line). The colored boxes on the graph, and above the images on the left-hand side, represent the different phases of motion (explained in C). **(C)** Kinetochore tracking revealed four distinct phases of motion as summarized in the cartoon on the left-hand side (described in detail in the main text). The bold lines in the theoretical graph on the right-hand side show the characteristic changes in distance to the different cellular locations (indicated in A) for the four phases. Phase 2 is characterized by rapid reduction in distance to the nearest spindle pole while Phase 4 by increase in distance to the spindle pole and reduction in distance to the midpoint and mid-plane. Color coding for each phase is indicated by the colored frames and boxes in the cartoon and the graph. **(D)** Time-lapse images show two representative cells passing through the early stages of mitosis (prometaphase). Cells were as described in B except they were treated with azBB. The features of the graphs on the right-hand side of the time-lapse sequences are as described in B.

We first analyzed how the positions of kinetochores changed over time in control cells (i.e. without azBB), following NEBD. Although such changes varied among individual kinetochores, we identified the following two common features for individual kinetochores (Figure 2B, C): First, after a few minutes following NEBD, kinetochores moved rapidly towards one of the two spindle poles, i.e. the distance to a spindle pole was rapidly shortened (Figure 2B, C; pale blue area). This poleward kinetochore motion started immediately after the initial kinetochore interaction with a MT extending from the spindle pole. The poleward kinetochore motion likely reflects the rapid kinetochore motion along the lateral side of a MT, as previously reported ^7,9^.

Second, after a few minutes following the completion of the poleward kinetochore motion, the kinetochore moved towards the mid-plane between spindle poles, i.e. the distance to the spindle pole was enlarged while the distance to the spindle mid-plane was shortened (Figure 2B, C; pale yellow area). This kinetochore motion seemed to reflect chromosome congression toward the spindle mid-plane ^12,13^. Such kinetochore motion continued for a few minutes and, once the kinetochore came close to the spindle mid-plane, it started oscillatory motions around the metaphase plate.

Based on the above motions of individual kinetochores, we defined Phase 1–4 as follows (Figure 2C): Phase 1 was the period between NEBD and the start of the poleward kinetochore motion – the latter also matched the initial kinetochore-MT interaction (based on co-localization of CENP-B and MT signals). Phase 2 was the kinetochore motion towards a spindle pole (see above). Phase 3 was defined as the period between Phase 2 and the start of kinetochore motion towards the spindle mid-plane. Phase 3 was often short in control cells and, during Phase 3, no common characteristic kinetochore motions (e.g. directional motions) were observed. Phase 4 was defined as the kinetochore motion towards the spindle mid-plane (see above).

We then analyzed the change of kinetochore positions over time in azBB-treated cells (Figure 2D). After a few minutes following NEBD, rapid kinetochore motions towards a spindle pole were observed, which was defined as Phase 2, as in control cells. This allowed us to define Phase 1 (the period between NEBD and the start of Phase 2) and Phase 3 (its start was the end of Phase 2) in the same way for azBB-treated and control cells. However, for some kinetochores in azBB-treated cells, the kinetochore motion towards the spindle mid-plane (defined as Phase 4) occurred after a long delay (Figure 2D, top) or did not occur during observation (Figure 2D, bottom). Next, using this data, we conducted a detailed comparison of kinetochore motions during each phase between control and azBB-treated cells.

### The PANEM contraction facilitates the initial interaction of peripheral kinetochores with spindle MTs, but not their subsequent poleward motions

While the PANEM showed contraction after NEBD, it seemed that peripheral kinetochores, i.e. those localizing near the nucleus-cytoplasm boundary at NEBD, moved inward (see below). We hypothesized that the effect of PANEM contraction on kinetochore-MT interaction is greater for peripheral kinetochores than for the kinetochores localizing to the central part of the nucleus at NEBD (central kinetochores). Therefore, to study kinetochore-MT interaction, we analysed peripheral and central kinetochores separately: the former was defined as kinetochores localizing within 2 µm of the nucleus-cytoplasm boundary at NEBD while the latter were the other kinetochores (Figure 3A). Kinetochore motions in Phase 1-4 were defined as in the previous section and in the same way for peripheral kinetochores (Figure 2B, D; Figure 2 – figure supplement 1A) and central kinetochores (Figure 2 – figure supplement 1B, C).

**Figure 3:**
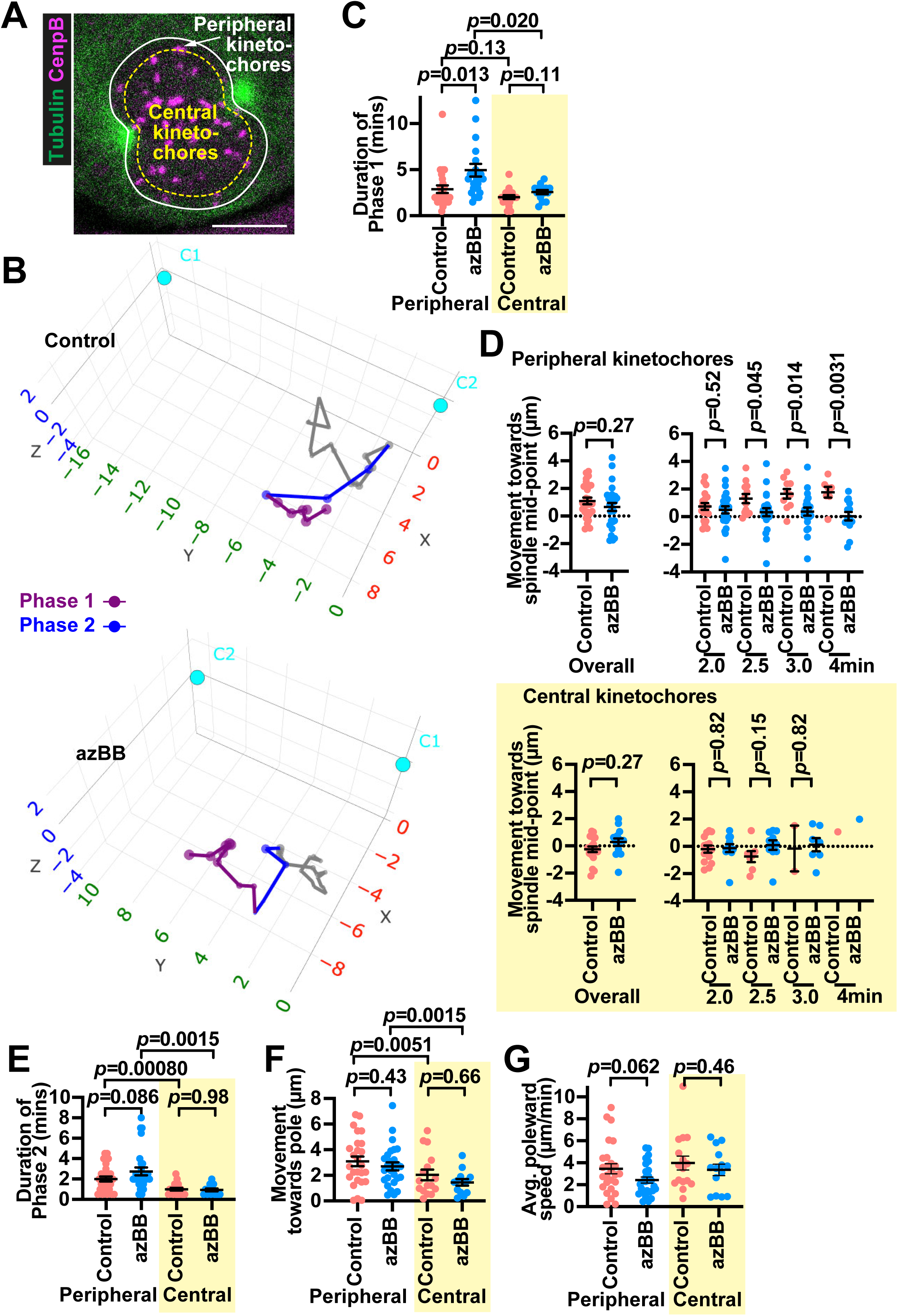
Reduced PANEM contraction affects motions of peripheral (but not central) kinetochores during Phase 1. **(A)** Image shows a cell in late prophase expressing CENPB-mCherry and GFP-⍰Tubulin. The white line indicates the position of the nuclear membrane. Kinetochores that fell within the yellow dotted line (positioned 2µm inside the nucleus) were considered central kinetochores, those between the two lines were considered peripheral kinetochores. There were a small number of non-kinetochore-derived mCherry signals, which localized outside the nucleus before NEBD and did not show any characteristic kinetochore motions, such as those towards a spindle pole and the spindle mid-plane, after NEBD. Scale bar is 10µm. **(B)** The motions of representative kinetochores are shown in 3D space over time in control cells (top) or azBB-treated cells (bottom). Kinetochores (colored dots) and spindle poles (turquoise circles) were tracked as in Figure 2 and positions were plotted relative to the nearest spindle pole (C2 in upper panel, C1 in lower panel) which was positioned at x=0, y=0, z=0. The position of the opposite spindle pole represents the average relative position of the pole during the time sequence. The colored lines of the kinetochore track represent Phase 1 (purple), Phase 2 (blue) as in Figure 2C. Phases 3 and 4 are shown in grey. The scales on all three axes are in µm. **(C)** Graph shows the duration of Phase 1 for individual kinetochores from control (red) or from azBB-treated (blue) cells. Kinetochores from the periphery are shown on the left-hand side and those from the center on the right-hand side with a yellow-colored box. The *p* values were obtained by t-test. 26, 27, 20 and 14 kinetochores (from 9, 9, 6 and 7 cells) were analyzed from left to right. Note that in the peripheral azBB-treated column one kinetochore took 17.5 minutes before entering phase 2 and is excluded from this graph. **(D)** Graph shows the distance moved towards the spindle mid-point during Phase 1 for individual kinetochores from control (red) or from azBB-treated (blue) cells. In each panel, the graph on the left-hand side shows the net change in distance of individual kinetochores towards the spindle mid-point, while the graphs on the right-hand side show the net change in distance during the indicated time period (relative to NEBD) for the subset of kinetochores that had not interacted with MTs during that period. The *p* values were obtained by t-test. **(E-G)** Graphs show the duration of Phase 2 (E), movement towards the nearest pole during Phase 2 (F) and the average poleward speed during Phase 2 (G) for individual kinetochores from control (red) or from azBB-treated (blue) cells. The source data for these analyses (coordinates of kinetochores and spindle poles) can be found in Figure 3 – source data 1 - 4.

Figure 3B shows examples of how the positions of peripheral kinetochores (after NEBD) changed over time, relative to the spindle pole (towards which the kinetochore moved during Phase 2), in control and azBB-treated cells. We first analyzed Phase 1 of the kinetochore motion. To investigate the effect of PANEM contraction on the initial kinetochore interaction with spindle MTs, we measured the duration of Phase 1. For peripheral kinetochores, Phase 1 was significantly longer in azBB-treated cells than in control cells (Figure 3C, left). By contrast, for central kinetochores, the duration of Phase 1 was not significantly different between azBB-treated cells and control cells (Figure 3C, right). This suggests that the PANEM contraction promotes the initial interactions of peripheral kinetochores, but not central kinetochores, with spindle MTs.

If PANEM contraction pushes peripheral chromosomes inward during Phase 1, the associated peripheral kinetochores may travel inward for a larger distance than central kinetochores. However, no significant difference was observed between azBB-treated and control cells, in the travel distance of peripheral kinetochores towards the mid-point of spindle poles during Phase 1 (Figure 3D, top, left). Central kinetochores also did not show a significant difference (Figure 3D, bottom, left). However, the travel distance measurement might be skewed since the period of Phase 1 for peripheral kinetochores was shorter in control cells than in azBB-treated cells (Figure 3C, left). Since, on average, Phase 1 finished earlier in control cells, peripheral kinetochores in these cells had less time to travel overall. To avoid this bias in our comparison, we measured the travel distance of sets of kinetochores before they interacted with spindle MTs, during several fixed time windows (2, 2.5, 3 or 4 minutes following NEBD). In this way, in each time window, only kinetochores that had not yet interacted with MTs were considered. In this analysis, the inward travel distance of peripheral kinetochores was significantly longer in control cells than in azBB-treated cells across the fixed time windows 2.5, 3 and 4 min (Figure 3D, top, right). By contrast, central kinetochores did not show such a difference (Figure 3D, bottom, right). These results suggest that the PANEM contraction moves peripheral kinetochores inward, but not central kinetochores, during Phase 1. We reason that such inward motions of peripheral kinetochores facilitate their initial interaction with spindle MTs, i.e. shorten the duration of Phase 1.

We next analyzed the kinetochore motions in control and azBB-treated cells during Phase 2. We found that the duration of Phase 2 was shorter for central kinetochores than for peripheral kinetochores (Figure 3E). However, there was no significant difference in the duration of Phase 2 between control and azBB-treated cells, for either the peripheral or central kinetochores (Figure 3E). We also analyzed the kinetochore travel distance towards the closest spindle pole during Phase 2, i.e. the difference in the distance to the spindle pole at the start and end of Phase 2. The kinetochore travel distance was shorter for central kinetochores than for peripheral kinetochores (Figure 3F). However, it showed no significant difference between control and azBB-treated cells, for peripheral or central kinetochores (Figure 3F). In addition, there was no significant difference in the average speed of the poleward motions between control and azBB-treated cells, for peripheral or central kinetochores (Figure 3G). These results suggest that, while the PANEM contraction reduces the time taken for kinetochores to interact with MTs, it does not affect the movement of kinetochores along MTs towards a spindle pole (Phase 2).

### The PANEM contraction helps peripheral kinetochores to start congression towards the spindle mid-plane, but does not promote their congressional motion itself

Figure 4A shows examples of how peripheral kinetochores subsequently changed their positions over time after NEBD, with Phase 3 and 4 motions highlighted. We analyzed Phase 3 of the kinetochore motions in control and azBB-treated cells. As expected, the start of Phase 3, relative to NEBD, was significantly delayed in azBB-treated cells (compared with control cells) for peripheral kinetochores, due to their extended Phase 1 (Figure 4B).

**Figure 4:**
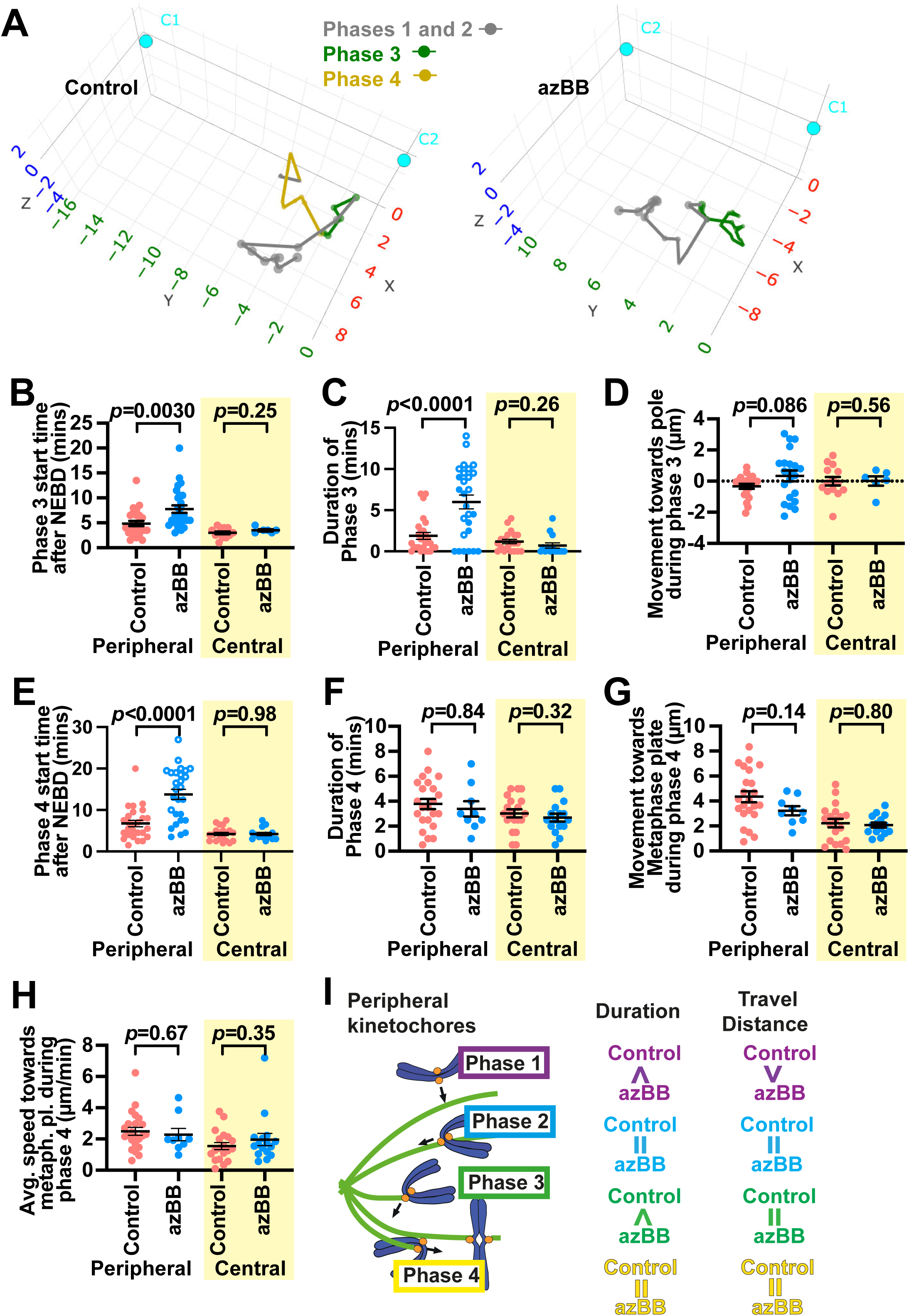
Reduced PANEM contraction leads to a delay in, or absence of, congressional movement of peripheral kinetochores. **(A)** The motions of representative kinetochores are shown in 3D space over time in control cells (left) or azBB-treated cells (right), as in Figure 3B. The colors of the kinetochore track represent Phase 3 (green), Phase 4 (yellow) as described in Figure 2C. The scales on all three axes are in µm. The same kinetochores as in Figure 3B are shown here. **(B–E)** Graphs shows the start time of Phase 3 relative to NEBD (B), duration of Phase 3 (C), net change in distance relative to the nearest spindle pole (difference between the start and end of Phase 3 [or the end of observation if congression did not start during observation]; D), and the start of Phase 4 relative to NEBD (or the end of observation if congression did not start during observation; E) for individual kinetochores from control (red) or from azBB-treated (blue) cells. The *p* values were obtained by t-test. 26, 27, 20 and 14 kinetochores (from 9, 9, 6 and 7 cells) were analyzed from left to right. Open circles indicate kinetochores that did not start congression (the end of Phase 3 was not defined). **(F–H)** Graphs show the duration of Phase 4 (F), net change in distance relative to the spindle mid-plane during Phase 4 (G) and the average speed during Phase 4 (H) for individual kinetochores from control (red) or from azBB-treated (blue) cells. The *p* values were obtained by t-test. 23, 9, 19 and 14 kinetochores (from cells corresponding to B-E) were analyzed from left to right. **(I)** Summary of comparison between control and azBB-treated cells for peripheral kinetochore motions during early mitosis (prometaphase). Similarities (equal sign) or differences (greater than or less than sign) are shown between control and azBB-treated cells. When a phase starts later, it is indicated by a ‘greater’ sign. Red asterisks indicate presumed direct effects of reduced PANEM contraction.

We next assessed the duration of Phase 3 in control and azBB-treated cells. In both conditions, the duration of Phase 3 was less than 5 min for all central kinetochores (Figure 4C). It was also less than 5 min for most peripheral kinetochores in control cells. However, for most peripheral kinetochores after azBB treatment, Phase 3 duration was longer than 5 min (Figure 4C). In many of them, we did not observe the end of Phase 3, i.e. the kinetochore motion towards the spindle mid-plane (Phase 4) did not start (Figure 4C, open circles). On the other hand, during Phase 3, there was no significant change in the kinetochore positions (relative to a spindle pole) in control or azBB-treated cells for peripheral or central kinetochores (Figure 4D), as we may expect from the definition of Phase 3 (i.e. the interval between Phase 2 and Phase 4). In summary, when the PANEM contraction was inhibited, Phase 3 was significantly extended for many peripheral kinetochores, but not for central kinetochores.

We then analysed the kinetochore motions during Phase 4, i.e. congression towards the spindle mid-plane (Figure 4A). The start of Phase 4 (relative to NEBD) was similar in control and azBB-treated cells for central kinetochores (Figure 4E), but it was significantly delayed for many peripheral kinetochores in azBB-treated cells (compared with control cells) (Figure 4E), which is explained by their extended Phase 1 and Phase 3. In contrast, once Phase 4 started, there was no significant difference in the duration of Phase 4 for peripheral or central kinetochores between control and azBB-treated cells (Figure 4F). There was also no significant difference in kinetochore travel distance or the average congression speed during Phase 4 between control and azBB-treated cells (Figure 4G, H). In short, in this analysis, a clear effect of inhibition of PANEM contraction was the failure (or delay) of several peripheral kinetochores to begin congression. However, if and once congression began, peripheral kinetochores (and central kinetochores) showed no significant change in motions of congression after the PANEM contraction was inhibited.

Summarizing the analyses of Phases 1-4 in the previous and current sections, the durations of Phase 1 and Phase 3 were extended for peripheral kinetochores, but not for central kinetochores, when PANEM contraction was inhibited with azBB (Figure 4I). The inward travel distance was shortened for peripheral kinetochores (but not for central kinetochores) during Phase 1 after azBB treatment (Figure 4I). Thus, we conclude that PANEM contraction facilitates peripheral kinetochores’ (but not central kinetochores’) initial interaction with spindle MTs as well as their start of congression towards the spindle mid-plane. On the other hand, the PANEM contraction affects neither kinetochores’ motion towards a spindle pole (following their initial MT interaction) nor their congressional motion.

Next, we wanted to specifically analyse kinetochores of polar regions, many of which, up to now had been grouped as part of the group at the cell periphery. Before moving on to these analyses, we wanted to test whether the conclusions we made above held for non-polar, peripheral kinetochores (polar kinetochores excluded). Since polar regions were generally smaller than non-polar regions, most of our selected peripheral kinetochores were in non-polar regions (while all selected central kinetochores were in non-polar regions). In any case, we repeated our analyses of Phases 1-4 only for the peripheral kinetochores specifically in non-polar regions (Figure 4 – figure supplement 1). The results for non-polar peripheral kinetochores were still essentially the same as those obtained for all peripheral kinetochores shown in Figures 3 and 4.

### The PANEM contraction repositions chromosomes from polar regions to ensure timely congression towards the spindle mid-plane

We next analyzed the motion of kinetochores, which were localized in polar regions at NEBD (polar kinetochores) (Figure 5A). For many polar kinetochores, it was hard to discriminate Phase 1 and Phase 2 because the direction of kinetochore movement was similar between the two phases. Therefore, our analysis focused on the transition from Phase 3 to Phase 4, i.e. the start of chromosome congression towards the spindle mid-plane. In previous studies, the start of chromosome congression was identified as the most critical regulatory step to avoid the missegregation of chromosomes localizing at polar regions ^18,19^. Meanwhile, it was also reported that congression of polar kinetochores to the mid-plane was facilitated by the separation of centrosomes and spindle elongation because these motions help to pivot captured polar kinetochores (those interacting with a MT emanating from one spindle pole) into the space between the two spindle poles ^23^. To measure the specific effect of PANEM contraction on polar chromosomes, we investigated cells where the start of chromosome congression should be minimally affected by spindle elongation. This was possible because, in U2OS *cdk1-as* cells (that we used in this study), it had been previously shown that arrest and release at the G2-M boundary leads to most cells having a fully (or almost fully) elongated spindle at NEBD ^21^. Among the cells selected in our experiments here, after NEBD, the spindle lengths were observed to gradually shorten, which occurred similarly between control and azBB-treated cells (Figure 1 – figure supplement 6).

**Figure 5:**
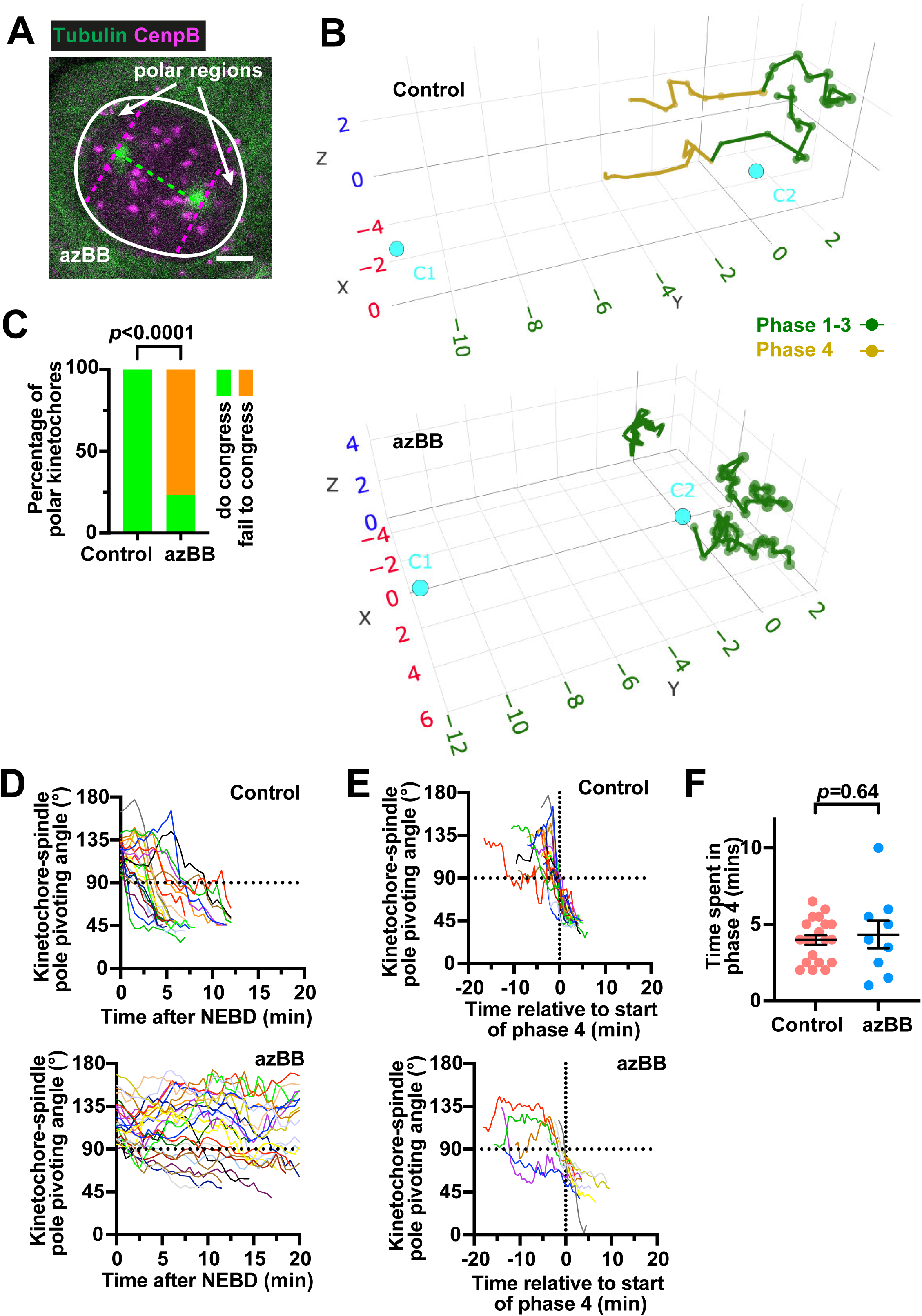
PANEM contraction is important to reposition kinetochores in polar regions at NEBD for efficient congression. **(A)** Image of a cell in late prophase expressing CENPB-mCherry and GFP-⍰Tubulin. The white line indicates the position of the nuclear membrane. The dotted green line is the line connecting the spindle poles. The magenta dotted lines are those on one spindle pole and perpendicular to the green dotted line. The polar regions are defined as the nuclear regions behind the pink dotted lines. Scale bar is 5µm. **(B)** The motions of representative kinetochores are shown in 3D space over time in control cells (top) or azBB-treated cells (bottom), as in Figure 3B. The colors of the kinetochore track represent Phase 1-3 (green), Phase 4 (yellow) as described in Figure 2C. The scales on all three axes are in µm. **(C)** Graph shows the percentage of polar kinetochores (at NEBD) that congressed in control (left) or azBB-treated (right) cells before the end of the time lapse sequence, or before tracking was no longer possible. The *p* values were obtained by Fisher’s exact test. Number of polar kinetochores was 23 and 22 from control and azBB-treated cells, respectively. **(D)** Plots show changes in pivot angles (defined as in Figure 2A) of polar kinetochores (at NEBD) over time after NEBD (time 0), from control cells (upper panel) and azBB-treated cells (lower panel). Individual colored lines indicate individual kinetochores. The dotted line indicates the angle at which a polar kinetochore (>90°) passed into the region between the poles (central region) (<90°). The number of polar kinetochores analyzed was 23 and 25 from 5 control and 2 azBB-treated cells, respectively. Figure 5 - figure supplement 2 shows the analyses of 35 polar kinetochores from 3 individual azBB-treated cells – the data only from the first two azBB-treated cells are shown in D to avoid overcrowding in the graph. **(E)** Plots show changes in pivot angles of polar kinetochores (at NEBD) over time, as in D but aligned according to start of congressional motion (time 0). Number of polar kinetochores analyzed was 23 and 10 from 5 control and 3 azBB-treated cells, respectively. For azBB-treated cells, plots show only the polar kinetochores that subsequently exhibited congression. **(F)** Graph shows the duration of Phase 4 for polar kinetochores (at NEBD) from control (red) or from azBB-treated (blue) cells. The *p* values were obtained by t-test. Number of polar kinetochores was 20 and 9 from 5 control and 3 azBB-treated cells, respectively. For azBB-treated cells, the graph includes only the polar kinetochores that subsequently showed congression. The source data for these analyses (coordinates of kinetochores and spindle poles) can be found in Figure 5 – source data 1 and 2.

Figure 5B shows examples of how polar kinetochores subsequently changed their positions over time after NEBD, relative to the nearest spindle pole, in control and azBB-treated cells. After identifying polar kinetochores at NEBD, we addressed whether they successfully congressed to the spindle mid-plane at later time points. While all polar kinetochores successfully congressed in control cells, 77% (23/30) of polar kinetochores failed to show congression in azBB-treated cells (Figure 5C). We then analyzed the angle between the spindle axis and the line from the kinetochore to the nearest spindle pole (Figure 2A, pivot angle): if a kinetochore was in the polar region, the angle was > 90°, and if in the region between the poles, it was < 90°. In control cells, most of the kinetochores moved from the polar region to the region between the poles (i.e. their angles became < 90°) within 10 min of NEBD (Figure 5D, control). After movement to the region between poles, congression (Phase 4) usually started soon after (Figure 5E, control). In contrast, in azBB-treated cells, the majority of polar kinetochores stayed in polar regions for 20 min (or more) after NEBD (Figure 5D, azBB, Figure 5 – figure supplement 1). A few kinetochores moved from the polar region to the region between the poles in azBB-treated cells, and in most such cases, similar to kinetochores from control cells, they showed congression soon afterwards (Figure 5E, azBB). Once congression started, kinetochores showed similar congression time, travel distance and speed in control and azBB-treated cells (Figures 5F, Figure 5 – figure supplement 2) as was the case for Phase 4 of peripheral non-polar kinetochores (Figure 4 – figure supplement 1J-L). These results suggest that the PANEM contraction facilitates movement of polar kinetochores out of polar regions and advances the onset of congression to the spindle mid-plane.

The PANEM contraction may achieve these effects by limiting the space of polar regions where polar chromosomes are located. To address this, we visualized both chromosomes and PANEM (in addition to the mitotic spindle) in control and azBB-treated cells with live-cell imaging (Figure 6A). We then quantified (1) the volume inside the PANEM at the back of a spindle pole (i.e. at the polar region) (Figure 6B) and (2) the chromosome volume present at the polar region (Figure 6C). Both volumes were rapidly reduced following NEBD in control cells, but the speed of reduction was diminished in azBB-treated cells (Figure 6B, C; Figure 6 – figure supplement 1). Thus, the polar region where polar chromosomes can be located was reduced by the PANEM contraction, which we reason facilitated their exit from the polar region.

**Figure 6:**
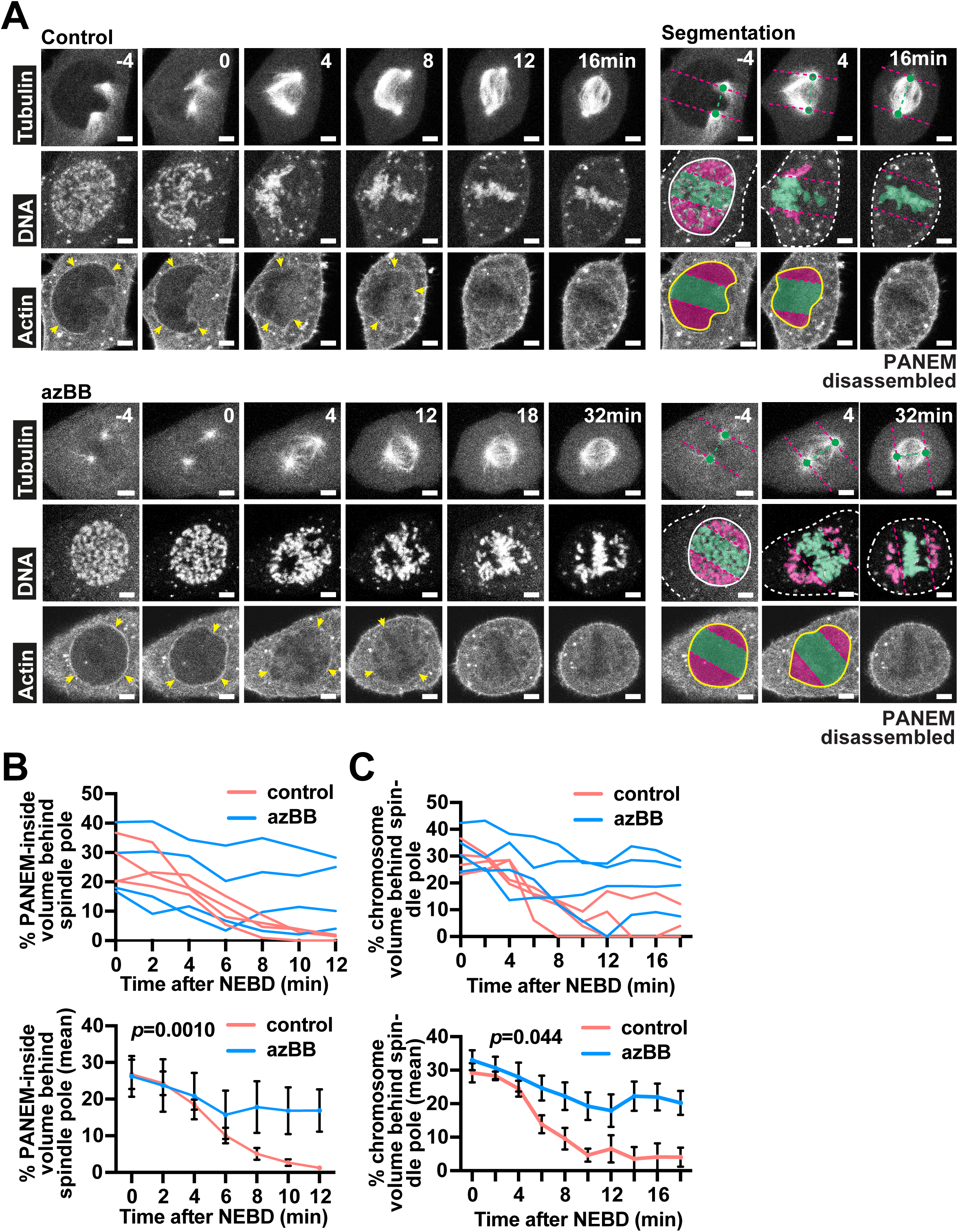
PANEM contraction leads to a rapid reduction of the PANEM-inside volume and chromosome volume at polar regions in early mitosis. **(A)** Time-lapse images show representative cells passing through the early stages of mitosis (prophase and prometaphase). A stable cell line expressing mCherry-LifeAct and GFP-⍰Tubulin, with chromosomes visualized by SiRDNA, were treated (or not) with azBB and were imaged every 2 minutes. Times shown were relative to NEBD. In the left-hand images the PANEM is indicated by yellow arrowheads. On the right-hand side, selected images have been reproduced to highlight segmentation. In the upper images, polar regions are designated by spindle poles (green dots) and their perpendicular planes (magenta dotted lines) (see Figure 5A). Chromosome (middle images) or PANEM (lower images) volumes behind or between the spindle poles are colored with magenta or green shading, respectively. In the middle images, solid white lines represent the cell nucleus (before NEBD) and white dotted lines represent the cell periphery. In the lower images, yellow lines represent the PANEM. Scale bars are 5µm. **(B and C)** Graphs show changes in PANEM-inside volume (B) and chromosome volume (C) behind the spindle poles as calculated for control (red lines) or azBB-treated (blue lines) cells. In the upper graph, the changes at individual polar regions are shown while in the lower graph, the change in mean is shown. The bars represent the SEM. The *p* value was obtained by t-test performed after regression analysis (see Figure 6 – figure supplement 1). In B and C, the same polar regions were analyzed.

### Evidence that the contractile PANEM directly pushes both polar chromosomes and non-polar peripheral chromosomes inward

The PANEM is formed on the outer surface of the NE during prophase ^20^. When the PANEM-inside volume is reduced by the PANEM contraction following NEBD, the PANEM (and the NE remnants underneath) may physically push chromosomes inwards at both polar regions and non-polar peripheral regions. To address this, we visualized both the PANEM and chromosomes (DNA) by live-cell microscopy (Figure 7A). PANEM was observed in all prophase cells (29 out of 29), prior to NEBD. We focused on the first 4.5 min after NEBD, during which Phase 1 was completed in most of the cells not treated with azBB (Figure 3C). We quantified the signals of PANEM and the DNA mass in selected orientations (Figure 7B) and plotted their intensities against the distance from the center of DNA mass over time (Figure 7C). Both PANEM and the outer edge of DNA moved inward over time, at both polar and non-polar regions (Figure 7C). During this process, PANEM was always positioned at the outer edge of DNA (Figure 7D, E). We obtained similar results in more cells (Figure 7 – figure supplement 1). These data provide evidence that the contractile PANEM pushes both polar chromosomes and non-polar peripheral chromosomes inward to reposition them.

**Figure 7:**
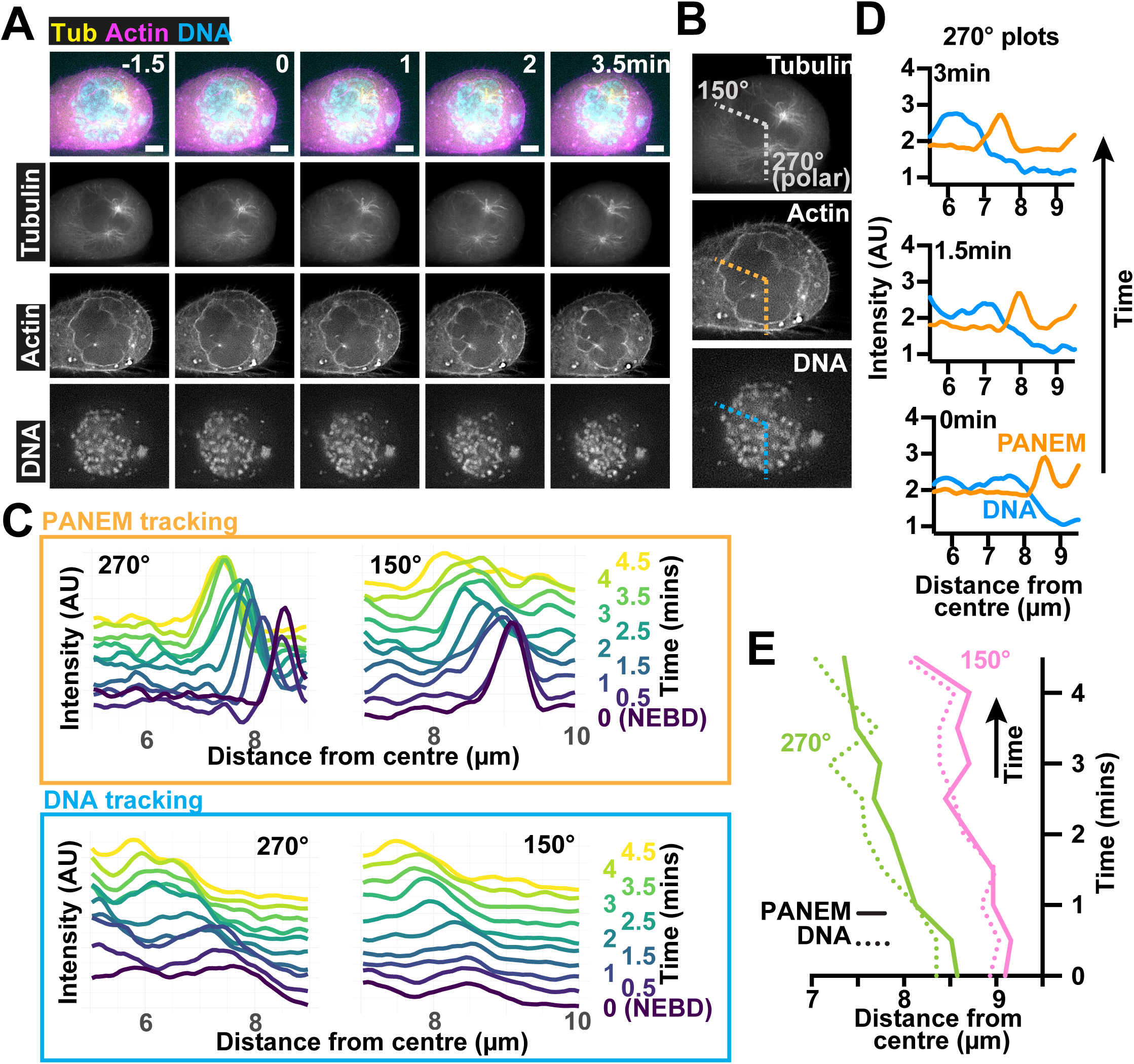
Evidence that the contractile PANEM directly pushes both polar chromosomes and non-polar peripheral chromosomes inward during early stages of mitosis. **(A)** Time-lapse images show a representative cell passing through the early stages of mitosis (prophase and prometaphase). A stable cell line expressing mCherry-LifeAct and GFP-⍰Tubulin, with chromosomes visualized by SYTO deep-red, were imaged every 30 seconds. NEBD is indicated at time 0. Scale bars 5µm. **(B)** Image 1.5min before NEBD from the time-lapse sequence in A to indicate the positions of line profiles. Dotted lines, plotted from the cell center (for determination see Materials and Methods), indicate the line profiles that pass the non-polar region (150°) and the polar region (270°) of the cell. **(C)** Graphs showing line profiles, offset in the y-axis according to time, for PANEM (upper panel; orange frame) or chromosomes (lower panel; blue frame) for the lines indicated in B. As time progresses peaks move to the left, which indicates movement closer towards the chromosome mass center. **(D)** Graphs show a time sequence of intensities calculated along the line profiles that pass the polar region of the cell shown in A. Time progresses upwards, and the colored lines indicate the intensities for Actin (PANEM; orange) or chromosomes (blue). **(E)** Graph shows the progression of the relationship between the PANEM peak and the chromosome front, over time, through the line profiles indicated in B.

### PANEM contraction helps eliminate polar chromosomes in multiple cell lines, while PANEM is absent in some chromosomally unstable cancer cells

While some cancer cell lines exhibit frequent chromosome missegregation, leading to numerical chromosomal instability (N-CIN+) and aneuploidy, other cell lines exhibit normal chromosome segregation and euploidy (N-CIN-) ^29^. In our previous study, in addition to U2OS cells, we found that PANEM forms in RPE1 cells (N-CIN-) but not in HeLa cells (N-CIN+) ^20^. We extended this analysis to additional human cancer cell lines (and also referred to a previous report ^21^), focusing on their N-CIN status. Intriguingly, PANEM formation was observed in all four N-CIN- cell lines, but was absent during prophase in 3 out of 5 N-CIN+ cell lines (Figure 8A, Figure 8 – figure supplement 1). This suggests that the absence of PANEM in some cancer cell lines may contribute to numerical chromosomal instability.

**Figure 8:**
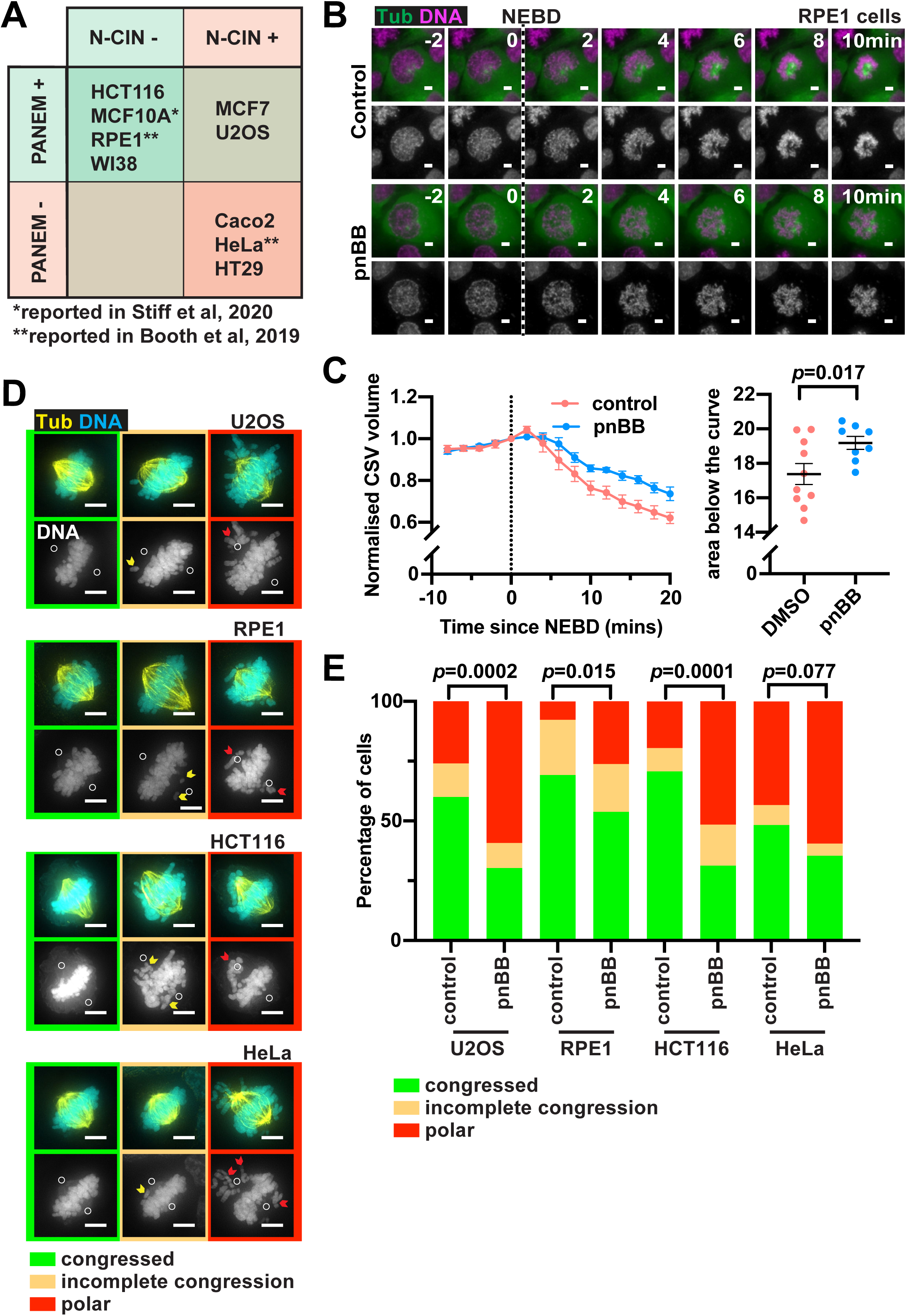
PANEM formation and function in non-cancer and cancer cell lines with and without numerical chromosomal instability. **(A)** Cell lines from this study (the U2OS cell line primarily used in this study and five cell lines in Figure 8 – figure supplement 1), from our previous study (HeLa and RPE1 ^20^), and from a study conducted by another group (MCF10A ^21^) were classified according to N-CIN+ or N-CIN-, i.e. with and without numerical chromosomal instability, respectively ^55^. Note that the PANEM status reported here for HCT116, RPE1 and U2OS cell lines were also confirmed by studies carried out independently of those from our lab ^21^. **(B)** Time-lapse images show a representative RPE1 cell passing through the early stages of mitosis (prophase and prometaphase). A stable cell line expressing GFP-⍰Tubulin, with chromosomes visualized by SYTO deep-red was imaged every 2 minutes after release from G2/M boundary. The timing of NEBD is indicated by the dotted line. Scale bar is 5µm. **(C)** The graph on the left shows the average change in normalized CSV for RPE1 cells imaged in B before and after NEBD (0 min) for cells treated with or without pnBB. The data from each cell was normalized to the volume at 0 minutes (immediately after NEBD) with standard error of mean (SEM) shown for each time point for each condition. The graph on the right plots the areas under the curves, measured in individual cells (which are shown in Figure 8 – figure supplement 2). The bars represent the mean and SEM. The number of cells for each group was 10 and 8 for control and pnBB, respectively. The *p* value was obtained using t-test. **(D)** Immunofluorescence of ⍰Tubulin and chromosomes (DAPI) in mitotic cells fixed 50 minutes after release from G2/M boundary. Different cell lines are shown from top to bottom, and different outcomes for each were observed, shown with coloured frames, from left to right. Those in green frames represent cells with successful alignment of all chromosomes; in orange some chromosomes had not completely aligned but were between the spindle poles (yellow arrow heads); in red some chromosomes had not aligned and remained behind the spindle poles (red arrow heads). In the lower image panels, showing DNA, for each cell line, the white circles represent the position of the spindle poles. Scale bars 5µm. **(E)** Quantification of chromosome alignment outcomes in different cell lines following treatment with DMSO (control) or pnBB for cells exemplified in D. Outcomes are shown in green, orange and red bars, as defined and using the same colors as in D. The *p* values were obtained using chi-square test for trends. The numbers of analyzed cells were 50, 76, 52, 80, 41, 64, 60 and 79 (left to right).

So far, we have studied the roles of PANEM contraction in U2OS cells. In U2OS cells, PANEM contraction rapidly reduced CSV following NEBD and eliminated polar chromosomes during prometaphase and metaphase ^20^ (Figure 1E-G). We next addressed whether PANEM contraction plays similar roles in other cell lines that form PANEM in early mitosis. First, we inhibited PANEM contraction with myosin-II inhibitor pnBB in RPE1 cells. Similar to what we observed in U2OS cells ^20^, pnBB treatment slowed CSV reduction following NEBD in RPE1 cells (Figure 8B, C; Figure 8 – figure supplement 2). In a second experiment, we arrested U2OS, RPE1, HCT116, and HeLa cells in late G2 with the Cdk1 inhibitor RO-3306, and subsequently released them into mitosis by washing out the inhibitor in the presence and absence of pnBB. With pnBB, the number of cells with polar chromosomes during metaphase increased in cell lines with PANEM (U2OS, RPE1, HCT116), but not significantly in the cell line without PANEM (HeLa) (Figure 8D, E). We conclude that the roles of PANEM in CSV reduction and in eliminating polar chromosomes are not limited to U2OS cells but are also found in other cell lines that form PANEM.

## Discussion

To ensure high-fidelity chromosome segregation, kinetochore-MT interactions must be efficiently and correctly established. This process is affected by the locations of chromosomes in cells: kinetochore-MT interactions occur less efficiently for chromosomes at the nuclear periphery or behind spindle poles (polar regions), resulting in a higher risk of mis-segregation during anaphase ^18,19^. The current study shows that contraction of the PANEM shortly after NEBD moves chromosomes, which were originally located at the nuclear periphery at NEBD, inward. This facilitates their kinetochores’ initial interaction with spindle microtubules (Phase 1 to 2, Figure 3) and also promotes the onset of their congression towards the spindle mid-plane (Phase 3 to 4, Figure 4). Notably, once these chromosomes initiate a poleward motion or congression (Phases 2, 4), the subsequent motions themselves are independent of PANEM contraction as might be predicted for MT-dependent motions (Figures 3 and 4). Moreover, the PANEM contraction reduces the volume of polar regions to help chromosomes escape from these regions, which allows initiation of their congression (Figures 5 and 6). The PANEM contraction seems to promote these processes by directly pushing chromosomes inward at the nuclear periphery and polar regions (Figure 7). Thus, the PANEM contraction relocates chromosomes from the regions where kinetochore-MT interactions are less efficient to regions where such interactions more readily occur (Figure 9). This ensures bi-orientation of sister chromatids and high-fidelity chromosome segregation.

**Figure 9:**
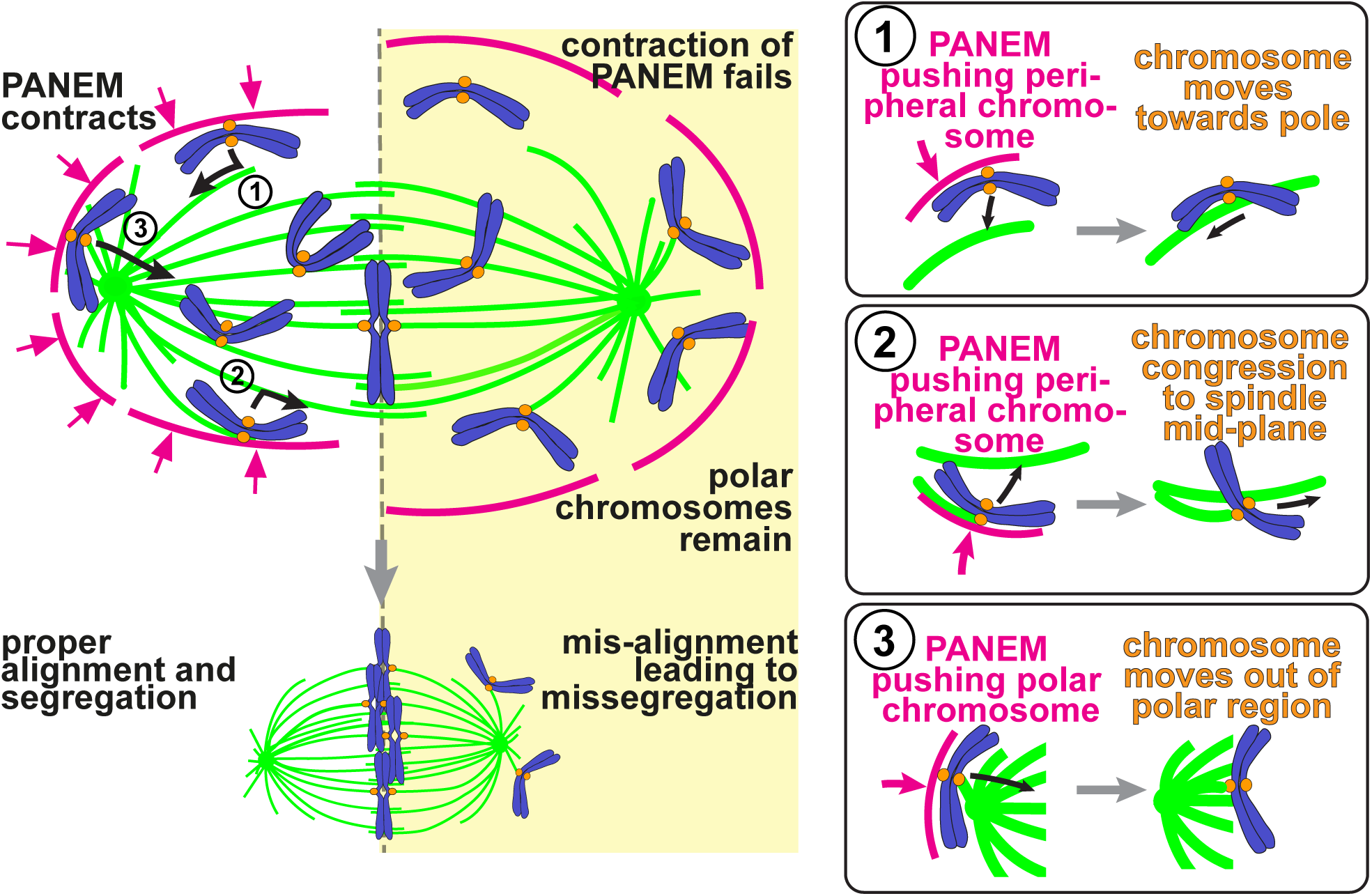
Model showing the effect of PANEM on peripheral/polar chromosomes. Left-hand side shows the model of how the PANEM contraction helps to prevent chromosome misalignment and subsequent missegregation (not shown) by pushing peripheral and polar chromosomes inward so that they can be efficiently captured on spindle MTs and subsequently transported to the spindle mid-plane where they become bioriented. With reduced PANEM contraction (yellow-coloured box), chromosomes often remain in polar regions. On the right-hand side, the cartoons framed in boxes show the three main effects of PANEM contraction: (1) initial capture of a kinetochore on a peripheral chromosome by a MT, emanating from one of the spindle poles (Phase 1), so that subsequent movement towards that pole starts efficiently; (2) second MT interaction of sister kinetochore on a peripheral chromosome (Phase 3) allowing the start of congression towards the spindle mid-plane; (3) relocation of a polar chromosome to the region between the spindle poles to facilitate productive MT interactions. Figure 9 – figure supplement 1 shows an alternative model of how PANEM contraction advances the onset of chromosome congression.

In our study, we exposed the whole nucleus to infrared light to activate azBB and inhibit myosin II on the PANEM. It was technically very difficult to activate azBB only at the NE regions where the PANEM is formed. One could argue that the defective kinetochore-MT interactions with azBB treatment, found in this study, might be due to inhibition of other actomyosin networks in the nucleus (e.g. present on centrosomes or on the spindle) ^30–33^. However, this is unlikely for the following reasons: (1) We did not detect any actin network inside the nucleus, on the spindle or between chromosomes in our study, at least, using the method and the cell line in the current study; (2) The inhibition of myosin II by azBB caused defective kinetochore-MT interactions for peripheral kinetochores, but not kinetochores located in central regions of the nucleus. If a possible actomyosin network in the nucleus, on the spindle or between chromosomes, were important for kinetochore-MT interaction in our system, we would expect defects for both peripheral and central kinetochores; and (3) While unconventional myosin-10 localizes at spindle poles and plays important roles in mitosis ^34,35^, it is unlikely to be affected in this study since, in contrast to myosin II, myosin-10 is not suppressed by blebbistatin ^33^, the parent myosin II inhibitor from which azBB was derived.

The PANEM contraction relocates chromosomes from polar regions and helps them initiate chromosome congression to the spindle mid-plane (Figure 5). A recent study suggested that centrosome separation and spindle elongation also play similar roles in helping chromosomes escape from polar regions and initiate congression ^23^. We speculate that the two mechanisms might work in parallel and likely cooperate. It has been shown that spindle elongation can occur with different timings between different cell lines and between different cells of the same cell line – in some, the spindle elongation occurs before NEBD, while in others after NEBD ^21,23,36^. Therefore, if spindle elongation is completed before NEBD but large polar regions remain when NEBD occurs (as we often observed in U2OS cells in the current study), the subsequent PANEM contraction would be important to eliminate the polar regions to help polar chromosomes escape from these regions. Alternatively, in cells where NEBD occurs before spindle elongation is completed, the spindle elongation and PANEM contraction could cooperate to move chromosomes out of polar regions. We intended to compare the effect of PANEM contraction in U2OS cells in which spindle elongation occurred before or after NEBD. However, we could not address this because spindle elongation usually occurred before NEBD in asynchronously growing U2OS cells, and elongation after NEBD was rare (and modest if observed; Figure 5 – figure supplement 3). In the future, it would be important to investigate how the two mechanisms – PANEM contraction and spindle elongation – cooperate to reduce polar chromosomes and under which conditions each mechanism is predominant (e.g. different cell lines, different timing of spindle elongation).

In the current study, we were able to identify kinetochore motions towards a spindle pole (Phase 2) and later, towards the spindle mid-plane (Phase 4), during prometaphase of U2OS cells (Figure 2). By contrast, a previous study, using RPE1 cells, suggested that kinetochores stayed near the spindle mid-plane and their motions towards a spindle pole and towards the spindle mid-plane were rather modest during prometaphase ^5^. This difference may be explained by the different timing of spindle elongation (the increase in distance between two spindle poles): This primarily occurred before NEBD in U2OS cells, while it mainly happened after NEBD in RPE1 cells ^5^.

It has been a widely accepted view that chromosomes congress to the spindle midplane before establishment of biorientation, whereby sister kinetochores interact with MTs from the opposite spindle poles ^12^. However, very recently, this view has been challenged by a new model where biorientation precedes congression and therefore promotes it ^25^. In any case, our results can be understood from both points of view. The first model (congression precedes biorientation) has been discussed in detail above. In the second model (where biorientation precedes congression), PANEM contraction could facilitate biorientation of non-polar peripheral chromosomes by constricting MTs from the spindle poles into a smaller region, thus focusing their extension towards chromosomes beyond the spindle midplane (Figure 9 – figure supplement 1). In this second situation, PANEM contraction would also facilitate biorientation of polar chromosomes by relocating them to non-polar regions with congression following ^18,22,25^.

Our results show that, during prophase, the PANEM is rapidly formed on the cytoplasmic side of the NE, relying on the LINC complex ^20^ (Figure 1A). Immediately after NEBD, the PANEM shows a rapid myosin-II-dependent contraction ^20^ (Figure 1C, D). Temporal regulation of the PANEM assembly and subsequent contraction should be crucial for the role of PANEM during prometaphase, which has been revealed in the current study. However, it is still unclear what triggers rapid PANEM assembly and contraction with such precise timing. PANEM assembly may be dependent on mitotic kinase activation of PANEM regulators (e.g. LINC complex ^37^, actin nucleators ^38^). PANEM contraction may be triggered by reduced tension on the NE (triggered by NEBD ^39^) or rapid accumulation of myosin-II on the perinuclear actin network ^20^. In the future, uncovering the molecular mechanisms responsible for such temporal regulation will help to obtain a complete picture of PANEM regulation and function in early mitosis.

Our previous study used a dominant-negative LINC construct (LINC-DN) to impair the formation of PANEM ^20^. LINC-DN attenuated the reduction of CSV soon after NEBD and increased the number of polar chromosomes ^20^; i.e. in this regard, the outcome was similar to azBB treatment in the current study. One might also expect that global actin polymerization inhibitors would inhibit the PANEM formation and show effects similar to LINC-DN. By contrast, it was reported that global actin polymerization inhibitors (e.g., cytochalasin D, latrunculin A) strongly affect mitotic rounding and cytokinesis but only modestly influence early chromosome movements ^23,40,41^. One possibility is that such differences may have arisen from different cell types – this could be important, especially given that some cells form the PANEM and others do not (Figure 8A). A second possibility is that cytokinesis, mitotic rounding and PANEM formation may rely on actin polymerization to different extents. For example, the same concentration of global actin polymerization inhibitors may affect cytokinesis, but may still allow PANEM formation to proceed without observable effects on early chromosome movements.

It has been demonstrated that the actin network cooperates with the spindle MTs to allow high-fidelity chromosome segregation. Well-known examples are the actin network along the cell cortex, which regulates the spindle orientation and positioning ^42,43^, and the actomyosin ring at the cell division site, which promotes cytokinesis ^44^. In addition, actin networks on/around the nucleus and within the spindle also facilitate chromosome segregation in mitosis and meiosis ^45^: For example, in starfish oocytes, the actin depolymerization underneath the NE remnants facilitates chromosome interaction with spindle MTs ^46,47^. In mammalian oocytes, actin dynamics on the spindle promote robust kinetochore-MT interactions and chromosome clustering to ensure chromosome segregation ^48,49^. In mouse preimplantation embryos, the actin network is formed between chromosomes and its contraction due to actin disassembly helps chromosome clustering and congression ^50^. In these systems, the actin network contributes to the proper positioning of chromosomes, which are widely scattered in oocytes and cells of the early embryo, to prepare for their subsequent segregation. In the present study, we describe a crucial role of a distinct actin network in repositioning chromosomes from unfavorable locations to facilitate their interaction with the mitotic spindle in human cells. These studies highlight the remarkable adaptability of actin networks in assisting the spindle MTs to safeguard faithful chromosome segregation during cell division of different cell types and across different species.

Finally, we investigated PANEM formation in N-CIN+ and N-CIN- cell lines. Intriguingly, PANEM formation was observed during prophase in all four N-CIN- cell lines, but was absent at this time in 3 out of 5 N-CIN+ cell lines (Figure 8A). These results raise two interesting possibilities. First, the absence of PANEM in some cancer cell lines may itself contribute to numerical chromosome instability. In the future, to test this possibility, a systematic study of a wider range of cancer cells and normal cells will be required. Second, cells without PANEM might have evolved compensatory mechanisms for efficiently establishing interactions between chromosomes and the spindle MTs (e.g. required for chromosome congression): For example, (1) the enhanced assembly rate of spindle MTs may facilitate kinetochore-MT interactions in N-CIN+ cancer cells ^51^; (2) chromosome biorientation may precede congression more frequently to promote the congression towards the spindle midplane ^25^; or (3) the balance between CENP-E, Dynein and chromokinesin’s activities may incline to greater chromosome-arm ejection forces towards the spindle midplane ^22^. If PANEM is widespread in normal tissues, then targeting these compensatory mechanisms could open opportunities for selectively killing PANEM-deficient cancer cells.

## Supporting information

Response to reviewers

## Acknowledgements

We thank Tanaka lab members, K. Weijer, A. Saurin and M. McLean for discussion; Dundee Imaging Facility for help with microscopy; and H. Hochegger, M. Lampson, J. Swedlow, J. Buerstedde and M. Davidson for reagents. This work was supported by the Wellcome Trust Investigator Award (219418/Z/19/Z), the Cunningham Trust PhD scholarship (CT19-06) and the Wellcome Trust Technology Platform (097945/B/11/Z). For open access, the authors will apply a Creative Commons Attribution (CC BY) license to any author-accepted manuscript version arising from this manuscript.

## Declaration of interests

One of the authors, A.J.R. Booth, is currently employed by IMSOL, providing service and maintenance of the DeltaVision Elite microscope used in this work. The microscope was also originally purchased through IMSOL.

## Author contributions

Conceptualization: T.U.T., N.S. and J.K.E.; methodology: N.S., J.K.E., Z.Y. and A.J.R.B.; investigation: N.S. and J.K.E.; formal analysis: N.S., J.K.E. and T.U.T., software: G.B. and J.K.E.; visualization: N.S. J.K.E. and T.U.T., writing – original draft: T.U.T., N.S, and J.K.E.; writing – review and editing: T.U.T. and J.K.E.: project administration, supervision and funding acquisition: T.U.T.

## Data availability

Coordinates of kinetochores and spindle poles, from which kinetochore motions relative to the mitotic spindle were analysed, are provided in the source data Figure 3 - source data 1–4 and Figure 5 – source data 1–2. Custom-made algorithms for this study are publicly shared at: https://doi.org/10.5281/zenodo.19822203 and https://github.com/graemeball/eLife_PANEM_scripts.

## Materials and Methods

### Cell Culture

The human cell line U2OS *cdk1-as* (male) was created by and obtained from the laboratory of Helfrid Hochegger ^52^. The human cell lines MCF7 (ATCC HTB-22; female), WI38 (ATCC CCL-75; female), HT29 (ATCC HTB-38; female), HCT116 (ATCC CCL-247; male), Caco2 (ATCC HTB-37; male) and hTERT RPE1 (referred to in this manuscript as RPE1; ATCC CRL-4000; female) were all obtained from American Type Culture Collection. U2OS *cdk1-as* cells and derivative cell lines were cultured at 37°C and 5% CO_2_ under humidified conditions in DMEM (with L-glutamine) (Thermo Fisher; 41966-052), 10% FBS (Thermo Fisher; 10270106), 100 U/ml penicillin and 100µg/ml streptomycin (Thermo Fisher; 15140-122). For live-cell microscopy, the above medium was replaced with Fluorobrite DMEM medium (Thermo Fisher; A18967-01) supplemented with 10% FBS, 2mM L-Glutamine (Thermo Fisher; 25030-024), 1mM Sodium Pyruvate (Lonza; 13-115E) and 25 mM Hepes (Thermo Fisher; 15630080). WI38 and MCF7 cells were cultured under the same conditions as above. Caco2 cells were grown under the same conditions as above except with medium supplemented with 20% FBS. HT29 and HCT116 cells were grown under the same conditions except using McCoys 5a Medium (Thermo Fisher; 26600080) supplemented with 10% FBS. RPE1 cells were grown under the same conditions except using DMEM-F12 medium (Thermo Fisher; 31331028) supplemented with 10% FBS and 100 U/ml penicillin and 100µg/ml streptomycin. Cell lines were regularly tested for mycoplasma contamination using kits from Lonza (LT07-118) or from Invivogen (rep-mys-10). No cell lines in this study were found to be contaminated with mycoplasma. Transfection of plasmids into U2OS *cdk1-as* cells or derivative cell lines was carried out using Fugene HD (Promega; E2311) according to manufacturer guidelines. Briefly, cells were transfected in single wells of a 6-well dish using 3µl Fugene HD and 1µg plasmid (3:1 ratio). The selection agents G418 (Formedium; G418S; 300µg/ml) or Blasticidin (Invivogen; ant-bl-1; 2µg/ml) were introduced 24-48 hours after transfection. Transfection of plasmids into RPE1 cells was carried out using the NEON Electroporation system (Thermo Fisher). For each transfection, 1 x 10^6^ cells were electroporated with up to 1.5µg of plasmid DNA according to manufacturer’s guidelines. Briefly, the cells were resuspended in 50µl of resuspension buffer and cells/DNA were electroporated sequentially in 10µl batches using the following conditions 1050V; 30ms width; 2 pulses. Cells were then placed immediately into 0.5ml fresh medium before being spun-down and resuspended in 2ml fresh medium in a single well of a 6-well dish. The selection agent G418 (Formedium; G418S; 500µg/ml) was introduced 24-48 hours after transfection.

U2OS *cdk1-as* (and derivative) cells expressing CDK1as were synchronized at the G2/M boundary using 1NMPP1 (Sigma; 529581-1MG). Briefly 0.2 x 10^6^ cells were seeded in 2 ml of medium in 3cm glass-bottomed microscope dishes (World Precision Instruments). 16 hours before imaging 1NMPP1 was added at a final concentration of 1µM and incubated for 12 to 16 hours. 1NMPP1 was then removed and cells were washed with 10 x 1ml of fresh medium to release cells into mitosis.

In the cell lines that did not contain the *cdk1-as* allele, enrichment of cells at the G2/M boundary was achieved using the CDK1 inhibitor RO-3306 (Selleck Chemicals; S7747). Briefly, 0.2 x 10^6^ cells were seeded in 2 ml of medium in 3cm glass-bottomed microscope dishes (World Precision Instruments). 6-8 hours before imaging, RO-3306 was added at a final concentration of 8µM and incubated for 6 to 8 hours. Medium containing RO-3306 was then removed and cells were washed with 10 x 1ml of fresh medium to release cells into mitosis.

The microtubule destabilizer Nocodazole (Sigma M-1404) was used at a concentration of 3.3µM and was added to the cells 1 hour before imaging. The Wee1 inhibitor MK-1775 (Selleck Chemicals; S1525) was added to cells with nocodazole or pnBB at a concentration of 0.5µM to facilitate release from G2/M arrest. The DNA stain SiR-DNA (Spirochrome/Tebubio; SC007) was used at a concentration of 100nM and was added to cells approximately 16 hours before imaging. The DNA stain SYTO-deep red (Thermo Fisher; S34900) was used at a concentration of 1µM and was added 2-3 hours before imaging. The myosin II inhibitor para-nitroBlebbistatin (pnBB; motorpharmacology/Tocris) was used at a concentration of 10-50µM and the photoactivatable Myosin II inhibitor azido-Blebbistatin (azBB; motorpharmacology) was used at a concentration of 5µM. Both of these were added to cells immediately after release from G2/M arrest (just before imaging).

### Plasmid construction

For stably expressing CENPB-mCherry in G418 resistant U2OS cells the plasmid pT3570 was generated. It is a derivative of pCENPB-mCherry (Addgene 45219; a gift from the laboratory of Michael Lampson) with the G418 resistance marker removed and replaced with a Blasticidin resistance marker from ploxBlastR ^53^.

### Cell line construction

The human cell line U2OS *cdk1-as* (mentioned above), whose endogenous CDK1 genes were disrupted and that expressed *Xenopus* CDK1as transgene, was used to generate different cell lines for microscopy experiments. A derivative of this cell line expressing GFP-⍰Tubulin under the control of the CMV promoter, designated TT215, was created by transfecting U2OS *cdk1-as* cells with the plasmid pGFP-⍰Tubulin (a gift from the laboratory of Jason Swedlow) and selection for G418 resistance. A second derivative of U2OS *cdk1-as* that expressed GFP-⍰Tubulin (as above) as well as H2B-mCherry, designated TT113, was created by co-transfecting cells with pGFP-⍰Tubulin (as above) and pH2B-mCherry (a gift from the laboratory of Jason Swedlow) and selection for G418 resistance. A third derivative of U2OS *cdk1-as* that expressed GFP-⍰Tubulin (as above) as well as mCherry-Lifeact-7 under the control of the CMV promoter, designated TT124, was created by co-transfecting cells with pGFP-⍰Tubulin (as above) and pmCherry-Lifeact-7 (Addgene 54491; a gift from the laboratory of Michael Davidson) and selection for G418 resistance. Finally, a derivative of TT215 (as above) expressing CENPB-mCherry under the control of the CMV promoter, designated TT230, was created by transfecting cells with the plasmid pT3570 (see above section) and selection for blasticidin resistance. A derivative of the RPE1 cell line expressing GFP-⍰Tubulin under the control of the CMV promoter, designated TT352, was created by transfecting RPE1 cells with the plasmid pGFP-⍰Tubulin (a gift from the laboratory of Jason Swedlow) and selection for G418 resistance. Stable cell lines used in this study were verified by Eurofin authentication service, using STR profiling.

### Microscope set-up, image acquisition and deconvolution

For microscope imaging that did not require the infra-red irradiation, time-lapse, live cell images were collected at 37°C with 5% CO_2_ while fixed cell images were collected at 25°C using a DeltaVision ELITE microscope (Applied Precision). We used an apochromatic 100x objective lens (Olympus; numerical aperture: 1.40) or 60x objective lens (Olympus; numerical aperture: 1.42) to minimize longitudinal chromatic aberration. We also routinely checked lateral and longitudinal chromatic aberration using 100-nm multi-color beads. We did not detect any chromatic aberration between the colors observed in the current study. For signal detection we used a sCMOS camera (PCO Edge) or a Cascade 1K EMCCD camera.

To visualize cells in early mitosis, U2OS *cdk1-as* cells and derivatives were arrested at the G2/M phase boundary, using 1NMPP1 before release into mitosis (see above). Alternatively, RPE1 (and derivatives), HCT116, U2OS (no cdk1as) and HeLa were arrested at the G2/M phase boundary, using RO-3306, before release into mitosis (see above). For visualising the chromosomes and ⍰Tubulin in TT113 cells, mCherry and GFP signals were discriminated using the dichroic DAPI/FITC/mCherry/Cy5 (52-852112-001 from API). Timelapse images, in each of these channels, were acquired through 10 z-sections separated by 2µm every 2 minutes for about 3 hours using 2x2 binning. For visualising chromosomes, actin and ⍰Tubulin in TT124 cells, SYTO-Deep red, mCherry and GFP signals were discriminated using the dichroic DAPI/FITC/mCherry/Cy5 (52-852112-001 from API). For tracking early stages of PANEM contraction, timelapse images, in each of these channels, were acquired through 20 z-sections separated by 0.5µm every 30 seconds for about 1 hour using 1x1 binning. For tracking PANEM formation and dissolution, timelapse images, in each of these channels, were acquired through 14 z-sections separated by 0.75µm every 1.5 minutes for about 1 hour using 1x1 binning. For visualising chromosomes and ⍰Tubulin in TT352 cells, SYTO-Deep red and GFP signals were discriminated using the dichroic DAPI/FITC/mCherry/Cy5 (52-852112-001 from API). To track changes to CSV in such cells as they entered mitosis, timelapse images, in each of these channels, were acquired through 12 z-sections separated by 1.5µm every 5 minutes using 4x4 binning and a 60x objective.

To visualize chromosomes and actin in PFA fixed cells (various cell lines) Hoechst and phalloidin DyLight 650 signals were discriminated using the dichroic DAPI/FITC/mCherry/Cy5 (52-852112-001 from API). To visualize ⍰Tubulin and DAPI in PFA fixed immunostained cells, DAPI signal and the secondary antibody label Alexa488 were discriminated using the dichroic DAPI/FITC/TRITC/Cy5 (52-852111-001 from API). Images were acquired through 25 z-sections of 0.5µm.

After acquisition, all images were deconvolved before analysis using softWoRx software with enhanced ratio and 10 iterations. Analysis of individual cells was performed using Imaris software (Bitplane) or ImageJ ^54^.

For experiments that used azBB treatment that required photoactivation a Zeiss confocal 710 system with 63x objective lens (NA 1.4) was used. For photoactivation of azBB a coherent Chameleon multiphoton laser was used. AzBB photoactivation experiments were carried out in cells either expressing GFP-⍰Tubulin and CenpB-mCherry or GFP-⍰Tubulin and mCherry-LifeAct. In both cell lines chromosomes were stained with SiRDNA. To focus on early mitotic cells, before NEBD, prophase cells were identified and selected as those with an intact nucleus displaying early signs of chromosome condensation (note that they were confirmed later in the experiment as mitotic when NEBD was observed). After identification of a suitable candidate, a zoom factor of approximately 6x was applied to include only the nucleus of the selected cell in the field of view. A reference image was taken to visualise ⍰Tubulin, kinetochores and chromosomes or ⍰Tubulin, actin and chromosomes by scanning with 488nm, 543nm and 633nm lasers in three z-sections 2µm apart for this xy region. At the same time the nuclear region was irradiated with an 860nm Chameleon multiphoton laser (0.2% power, pixel dwell time 6.64µsec) to activate azBB to allow its covalent binding to Myosin II. For tracking kinetochore motions, timelapse images were then acquired of the selected cell by scanning with only 488nm, 543nm and 633nm lasers in 11-15 z-sections 1.0µm apart with a pixel size of 0.085µm every 30 seconds for approximately 20-30 minutes. For measuring volume behind the spindle poles timelapse images were then acquired of the selected cell by scanning with only 488nm, 543nm and 633nm lasers in 11-15 z-sections 1.7µm apart with a pixel size of 0.128µm every 2 minutes for approximately 40-60 minutes.

After acquisition, analysis of individual cells was performed using Imaris software (Bitplane). Unless otherwise stated, for presenting images in figures, Actin was shown as a single z slice at the sharpest focal plane (where the PANEM contour was the clearest). For DNA and ⍰Tubulin, images were of several projected z slices to convey the features of the whole nucleus or cell.

### Cell fixation for visualization of chromosomes and actin or **⍰**Tubulin

For fixed-cell visualization, cells were grown in 3cm fluorodishes (WPI) and washed once with 2ml phosphate-buffered saline (PBS) which was then replace with 2ml 4% paraformaldehyde (PFA) in PBS for 10 minutes. The cells were then washed with 2ml PBS three times. PFA fixed cells were permeabilized by treatment with 2ml room temperature PBS containing 0.5% Triton for 10 minutes. The cells were then washed with 2ml PBS twice. For visualisation of chromosomes and actin, these cells were incubated with 0.5µg/ml Hoechst 33342 (Sigma-Aldrich 14533) and 0.002 Units/µl phalloidin DyLight650 (Cell Signalling Technology; 12956) in 2ml of PBS with 5% bovine serum albumin (BSA) overnight at 4°C. Finally, cells were washed with 2ml PBS with 5% BSA then covered with 2ml PBS and imaged immediately.

For visualisation of chromosomes and ⍰Tubulin, immunofluorescence was carried out. After fixation (as described above) these cells were incubated with PBS containing 3% BSA at 4°C for at least 2 hours. Antibody against ⍰Tubulin (Merck; mab1864) was diluted 1:1,000 in PBS containing 3% BSA and incubated with fixed cells overnight at 4°C. The cells were then washed with 2ml PBS containing 0.1% Triton three times before incubation with the secondary antibody Donkey anti-Rat Alexa488 (Invitrogen; A21208) that was diluted 1:1000 in PBS containing 3% BSA overnight in the dark at 4°C. The cells were then washed with 2ml PBS containing 0.1% Triton three times before 20µl Prolong Gold Antifade containing DAPI (Thermo Fisher; P36935) was mounted on cells, which were sealed with coverslips.

### Measuring chromosome scattering volume

Measurement of chromosome scattering volume (CSV) was carried out by a semi-automatic detection of chromosomes using Imaris (Version 8.0) software Surface tool, as described previously ^20^. Contours of chromosomes on each acquired Z-section were selected, excluding any background non-chromosome signal. Imaris then generated 3D objects, representing chromosomes surface for each time point, from the selected contours. An extension script, created for Imaris in MatLab, was used to calculate the minimum polyhedron to envelope the 3D surface (also known as a convex hull), the volume of which is the CSV (see also Figure 1B).

### Measuring changes in the volume inside PANEM

The analysis of the volume inside PANEM was carried out using Imaris (Version 8.0) software surface tool. Contours of the PANEM were drawn in each z-stack where the PANEM was visible as a full ring. Stacks where PANEM was indistinguishable from the actin network at the cell cortex were excluded. If the PANEM ring was detected in multiple z-stacks in a single timepoint, an average of the volume measurement in the z-stacks was taken to analyze the network contraction.

### Tracking and analysis of kinetochore motions

Cells were imaged with high temporal and spatial resolution, allowing tracking of kinetochores through time (see above). Tracking was carried out by automatic identification of kinetochores using Imaris software (Version 9.5), with user supervision and occasional correction. Using this feature, x, y, z coordinates of the center of mass of the kinetochores and centrosomes, visualized throughout prometaphase, were determined.

To understand the motions of the individual kinetochores through time, using the xyz coordinates for the different objects, the distance between the kinetochore and 3 fixed cellular points, were calculated (see also Figure 2A): (1) the spindle pole towards which the kinetochore moved after NEBD (usually the spindle pole that was closest to the kinetochore at NEBD); (2) the mid-point between the two spindle poles; (3) the shortest distance to the cell mid-plane (or metaphase plate). By setting these reference points on the spindle, we were able to analyze kinetochore motions without being affected by the motions of the spindle.

To easily visualise and clearly represent these kinetochore motions in relation to the spindle poles, a further round of processing was carried out using a python script. In brief, this script transformed the xyz coordinates of the spindle poles of a given sequence so that at each time point the closest spindle to the kinetochore at NEBD will locate at x=0, y=0, z=0, in addition the other spindle pole will always lie at x=0, z=0 (with variable y). According to how much the spindle pole positions had rotated or shifted, the kinetochore coordinates at each timepoint were transformed accordingly such that spindle pole motions were cancelled out. R studio was used to display these corrected data on a three-dimensional plot.

We assigned phases of kinetochore motions as follows: To define Phase 2, we looked at the changes in distances of the kinetochore from the nearest centrosome. When the distance dropped in 5 consecutive timepoints (or distance dropped more than 2 µm over the course of up to 5 timepoints), we recognised this as part of Phase 2. The start was then defined as the first moment the drop in distance began (after a period of no overall change, or of increasing distance change). In some cases, when distance to the nearest pole was already gradually dropping, we assigned the start of Phase 2 at the point where there was 2 to 4-fold increase in rate, which was usually also concomitant with an increase of the kinetochores’ distance from the mid-plane. When these distances changes were not clear, where it was possible, we assigned the start of Phase 2 according to the first time point we observed the overlap of the kinetochore signal with microtubules and subsequent movement towards the pole. The end of Phase 2 was assigned when the distance between the kinetochore and the nearest pole stopped shortening or began to rise slightly.

For non-polar kinetochores, the start of Phase 4 was assigned by looking for the time where the distance to the mid-plane began to decrease; usually, this was concomitant with an increase in distance from the nearest pole. The end of Phase 4 was assigned as a time when the kinetochore started oscillation (i.e. a backward motion towards a spindle pole) around the spindle mid-plane. In some cases where the kinetochore became close to the mid-plane (< 2.5 µm), it was not possible to track it further due to kinetochore crowding around the spindle mid-plane – in such cases, the end of Phase 4 was assigned as the end of tracking. For polar kinetochores, the above definition of the start of Phase 4 did not suit because the distance to the mid-plane already began to decrease before they entered the non-polar region. We therefore first specified the end of Phase 4 in the same way as above. We then looked at the corrected 3D plots of each kinetochore track (described in Materials and Methods above) and we traced the kinetochore track backwards, from the end point. We judged the start of Phase 4 for polar kinetochores as the first point that the kinetochore changed direction onto its final trajectory, of typically 4 or more consecutive timepoints, towards the mid-plane (congressional motion) after the kinetochore had passed into the region between the spindle poles.

We selected kinetochores that can be individually tracked. If kinetochore tracking was difficult because of kinetochore crowding (except towards the end of Phase 4; see above), we did not choose this kinetochore for further analyses. We also did not include kinetochores close to spindle poles (< 4 µm) at NEBD in our analysis for the following two reasons: First, these kinetochores often did not show clear and rapid movements towards a spindle pole, which we used to define Phase 2. Second, although we referred to kinetochore co-localization with a MT signal for the start of Phase 2, this was difficult for kinetochores close to spindle poles because of a high density of MTs. With such selection, all selected kinetochores without azBB treatment (control) showed the poleward motion (Phase 2) and congression (Phase 4) in this order, though their extents were varied among kinetochores. All selected kinetochores with azBB treatment also showed the poleward motion (Phase 2), and some of them showed congression (Phase 4) after Phase 2. Then, Phase 1 and Phase 3 were defined as intervals between NEBD and Phase 2 and between Phase 2 and Phase 4, respectively. If no Phase 4 was observed with azBB, we judged that Phase 3 continued until the end of tracking.

### Measuring PANEM and chromosome volumes behind the spindle poles

To determine the percentage of the PANEM volume lying behind the spindle poles, and to track changes of the volume through time, the following method was developed in Imaris: First, the x, y, z coordinates of the center of mass of spindle pole positions were determined automatically for each timepoint. Second, the PANEM volume or the chromosome volumes were determined for each timepoint. The PANEM volume was defined using the surface tool. Contours of the PANEM were drawn in each z-stack where the PANEM was visible as a full ring, stacks where the PANEM was indistinguishable from the actin network at the cell cortex were excluded. Finally, the volumes in each z-stack were unified to create a single surface object. The chromosome volume was defined by automatic detection using the surface tool. Background signals that were outside the nucleus before NEBD were excluded. For later stages of mitosis, when chromosomes became visible as individual units, all the chromosome volumes were unified as one surface item.

After the relevant objects of the cell had been tracked an extension script, created for Imaris in MatLab, was used to cut the surface objects of the PANEM or chromosomes into three surface objects. The regions were defined by taking the line joining the spindle poles and creating perpendicular planes to this line at each pole. Surfaces were then cut, along these planes, into up to three objects. They were either between the poles (i.e. between these planes), or behind one of the poles (not between the planes). For PANEM volumes, whose surface shape was simple, the split volumes at each time point were obtained directly. For chromosome volumes, whose surface shape was more complex, the surface cut script gave a simplified chromosome surface output. To calculate the chromosome volumes in each region accurately a further round of chromosome surface tracking was performed, using the same parameters as used originally, using the individual cut surfaces as a mask.

While the above analysis was carried out in 3D (includes all z stacks) for representation of the images in Figure 6A, a projection of several z-stacks was shown for the spindle poles and a single z slice was shown for both Actin and DNA images. Note that only polar regions with >20% total chromosome volume behind the spindle at NEBD were analyzed so that potential effects on the volume reduction due to PANEM contraction can be readily detected if present. For statistical analyses, Prism was used to fit linear regression lines for the data from each pole. For the PANEM volumes, regression was calculated for the first 12 minutes. For the DNA volumes, regression was calculated for the first 20 minutes. If zero % volume was reached and persisted for 2 or more timepoints in these analyses, the remaining points after the first zero were removed before regression was calculated.

### Tracking chromosome and PANEM motions using line plot profiles

We used both Imaris and ImageJ to track chromosome and PANEM movements, over time, relative to the center of the nucleus. First, the chromosome volume and its central x, y, z coordinates were defined by automatic detection using the surface tool in Imaris at each timepoint. ImageJ was then used to plot lines, arranged radially, from the chromosome center coordinates for each channel and timepoint using these coordinates. Intensity values along the length of each line were obtained. In some cases, where the selected center point displayed a poorly focused PANEM, images corresponding to plus or minus 1 z-stack were used for quantification, and these were matched between channels. Subsequently, for each selected line, data were normalized by dividing by the smallest intensity value found on that line and fluorescence intensity data were smoothed by averaging across 5 consecutive data points. The PANEM peak was determined as the point of highest fluorescence intensity. The DNA front was defined as the point halfway between the peak (defined as the last point of increasing intensity following at least three consecutive points of increase) and trough (defined as the last point of decreasing intensity following at least three consecutive points of decrease). For representation of the images in Figure 7A a projection of several z-stacks was shown for the spindle poles while a single z slice was shown for both Actin and DNA images.

### Statistical analysis and presentation

Graphs and statistical analyses were created and performed using Graphpad Prism (versions 7-10) and R studio (version 2024.12). All experiments were carried out at least twice and found to give consistent results. Number of cells analyzed, or data points included and the statistical tests used for each analyses are indicated in the Figure legends. The null hypotheses in statistical tests were that the samples were collected randomly and independently from the same population. All *p* values were two-tailed, and the null hypotheses were reasonably discarded when *p* values were smaller than 0.05.

## Supplemental Figure Legends

**Figure 1 - figure supplement 1:**
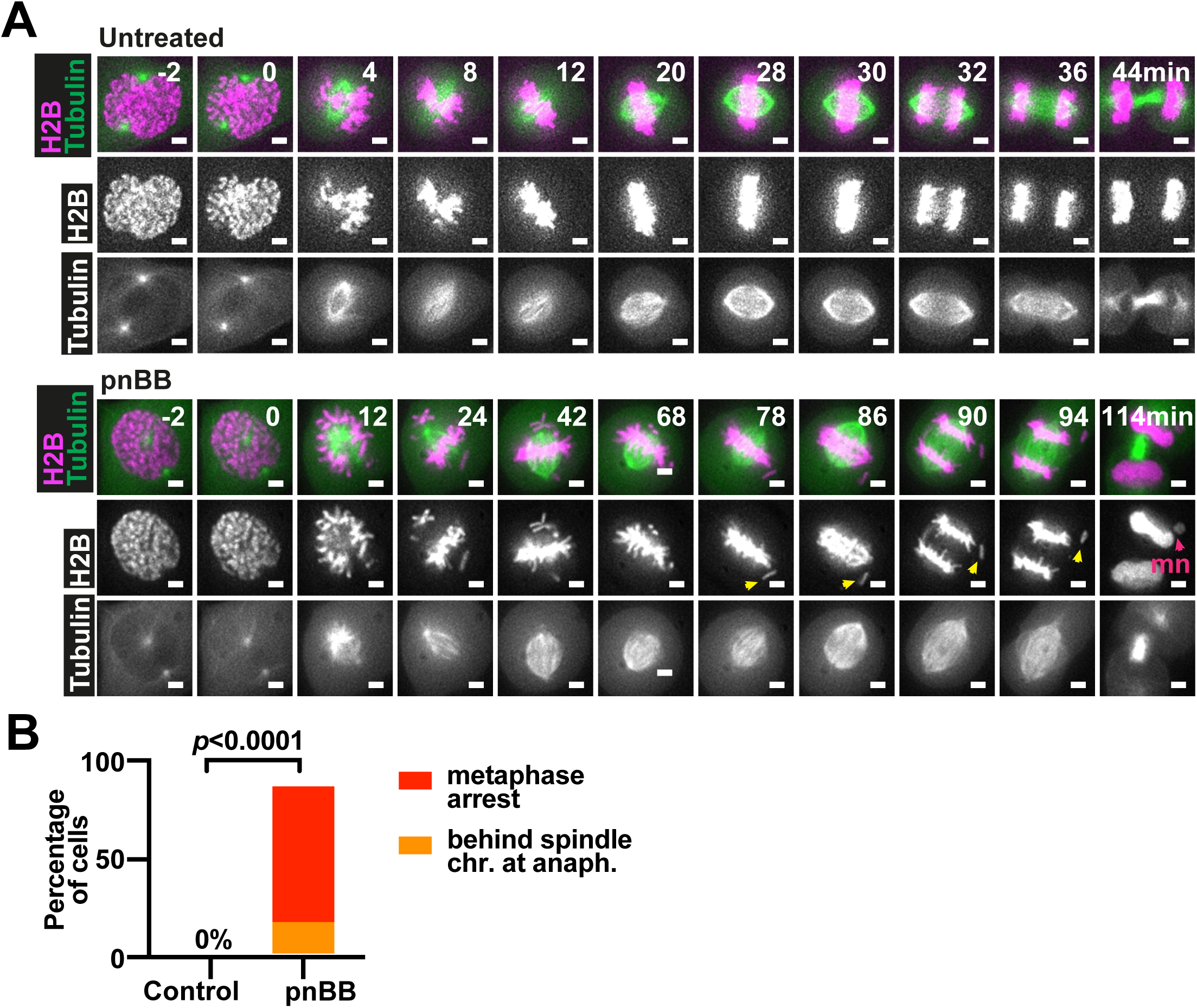
Chromosomes often show missegregation after PANEM contraction is inhibited. **(A)** Time-lapse images show representative cells passing through the different stages of mitosis (late prophase to telophase). A stable cell line expressing H2B-mCherry and GFP-⍰Tubulin was treated (or not) with pnBB and imaged every 2 minutes. Times shown were relative to NEBD. In the lower panel of images, a yellow arrowhead indicates a chromosome that was not aligned at the metaphase plate and was subsequently missegregated. At nuclear envelope reformation, this chromosome formed a micronucleus (mn; pink arrowhead). Scale bars are 5µm. **(B)** Graph shows the outcomes for mitotic cells as observed in time-lapse imaging exemplified in A. Cells that did not progress to anaphase after at least 2.5 hours after NEBD were considered arrested at metaphase (red). Many of these cells had apparent misaligned chromosomes locating behind spindle poles. An example of a cell that entered anaphase with chromosomes behind the spindle (orange) is shown in part A. Cells that progressed through anaphase normally (and did not show chromosome missegregation), exemplified in the upper panel of A, make up the remaining percentages of the graphs. Among the control cells, none of the cells became arrested at metaphase and all progressed to anaphase after proper alignment of chromosomes at metaphase. The *p* values were obtained by Fisher’s exact test. Number of cells analyzed was 15 and 19 from control and pnBB treated cells, respectively.

**Figure 1 – figure supplement 2:**
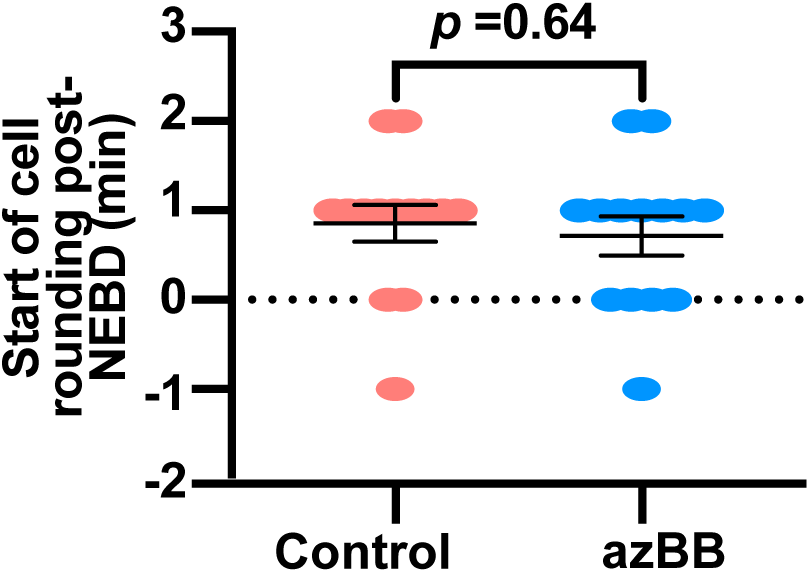
Inhibition of PANEM contraction does not lead to change in timing of mitotic cell rounding. Graph showing the timing of the start of cell rounding after NEBD for control (red) or azBB-treated (blue) cells. Cell rounding was determined as the first moment that the cell morphology changed as the cell started to detach from the growing surface. The bars show mean and SEM. The *p* values were obtained by t-test. Number of cells analyzed was 14 for both control and azBB treated cells.

**Figure 1 – figure supplement 3:**
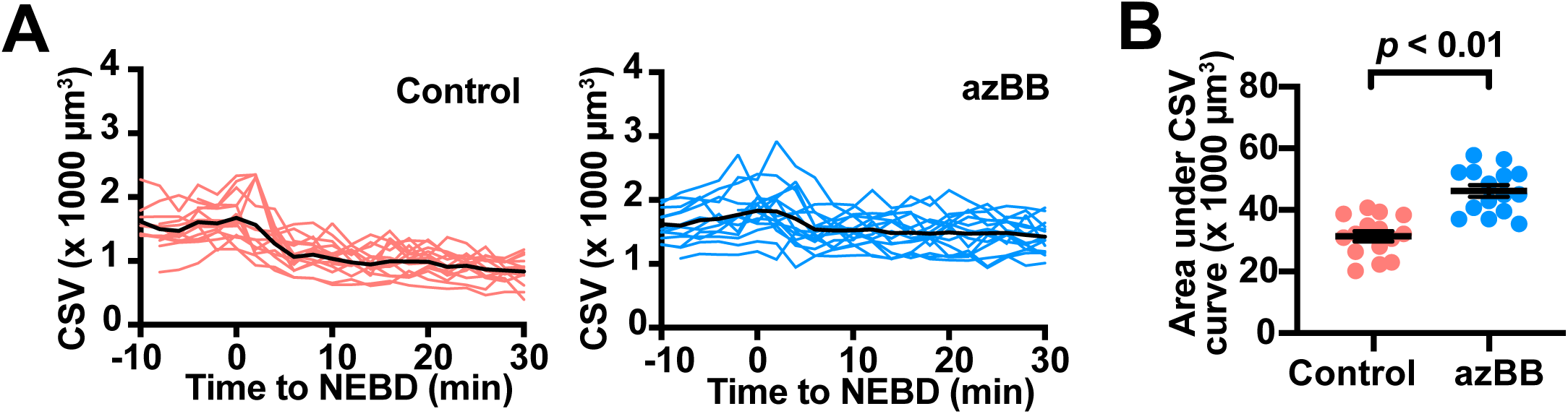
Inhibition of PANEM contraction leads to delay in reduction of CSV (data without normalization). **(A)** Graphs show chromosome scattering volume (CSV) before and after NEBD (0 min) for cells treated with or without azBB. This is from the same data shown in Figure 1D but before normalization. Each red or blue line represents the measurements from an individual cell while the heavy black lines represent the mean measurement across the time points. **(B)** Graph plots the areas under the curves for CSV (in A), measured in individual cells. The bars represent the mean and SEM. The *p* value was obtained using t-test.

**Figure 1 – figure supplement 4:**
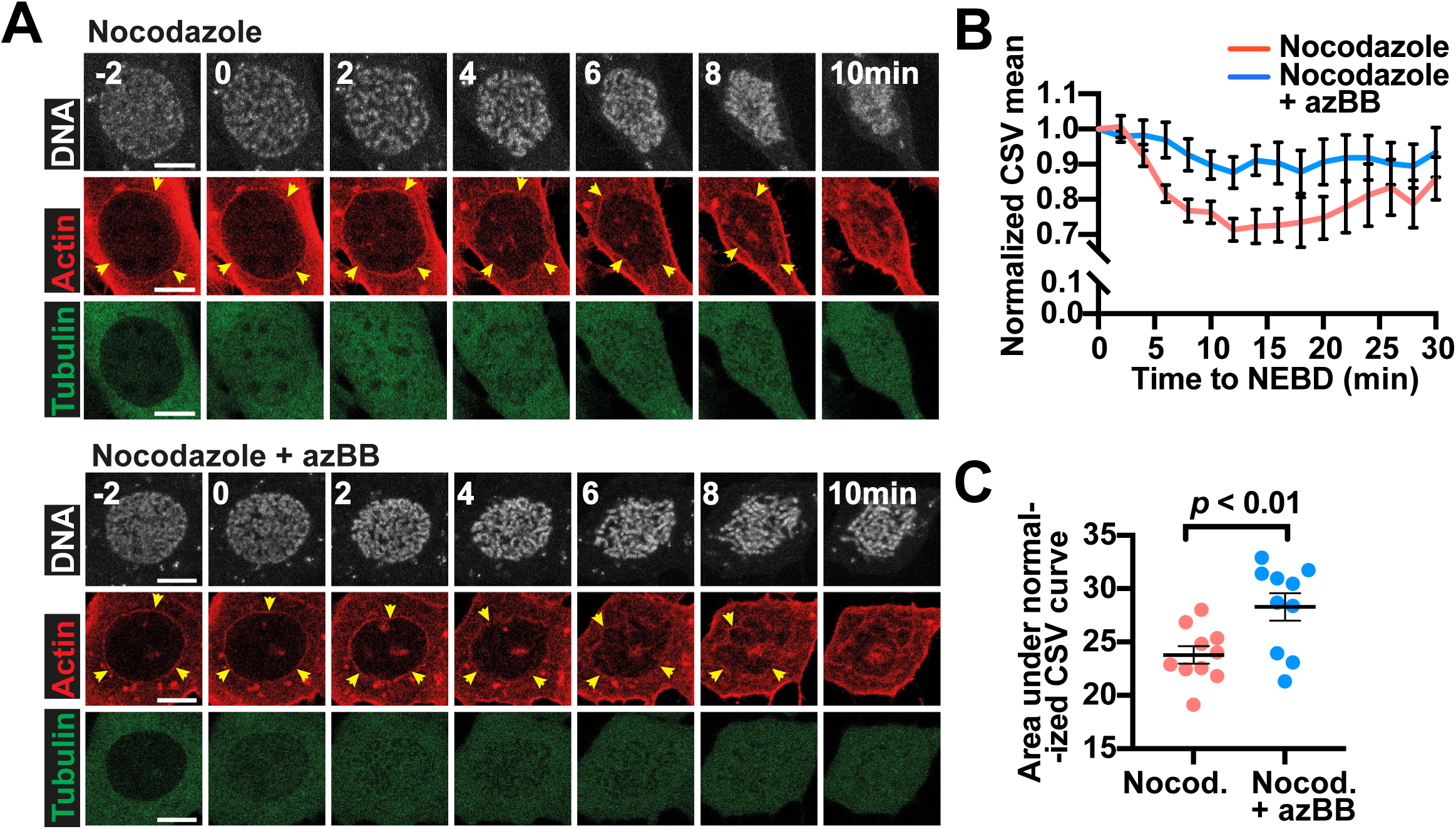
PANEM contraction and CSV reduction occurs in the absence of MTs. **(A)** Time-lapse images show representative cells passing through the early stages of mitosis (late prophase and prometaphase) following treatment with nocodazole and with or without azBB treatment. A stable cell line expressing mCherry-LifeAct and GFP-⍰Tubulin, with chromosomes visualized by SiRDNA, was imaged every 2 minutes. Times are relative to NEBD. The PANEM is indicated by yellow arrowheads. Scale bar is 10µm. **(B)** Graphs show chromosome scattering volume (CSV) after NEBD (0 min) for cells treated with nocodazole and with azBB (blue lines) or without azBB (red lines). Measurements were made from cells exemplified in A using methodology summarized in Figure 1B. The data from each cell was normalized to the volume at 0 minutes (immediately after NEBD). Here the mean, normalized values are presented with standard error of mean (SEM) shown for each timepoint for each condition. Normalized CSV mean, with nocodazole and without azBB (red line), slightly increased after 15 min, probably because chromosomes (not attached to spindle MTs) scattered again after the nuclear envelope remnants were lost. **(C)** The areas under the curves for CSV measurements in individual cells from data presented in B are shown here. The bars represent the mean and SEM. The *p* value obtained using t-test.

**Figure 1 – figure supplement 5:**
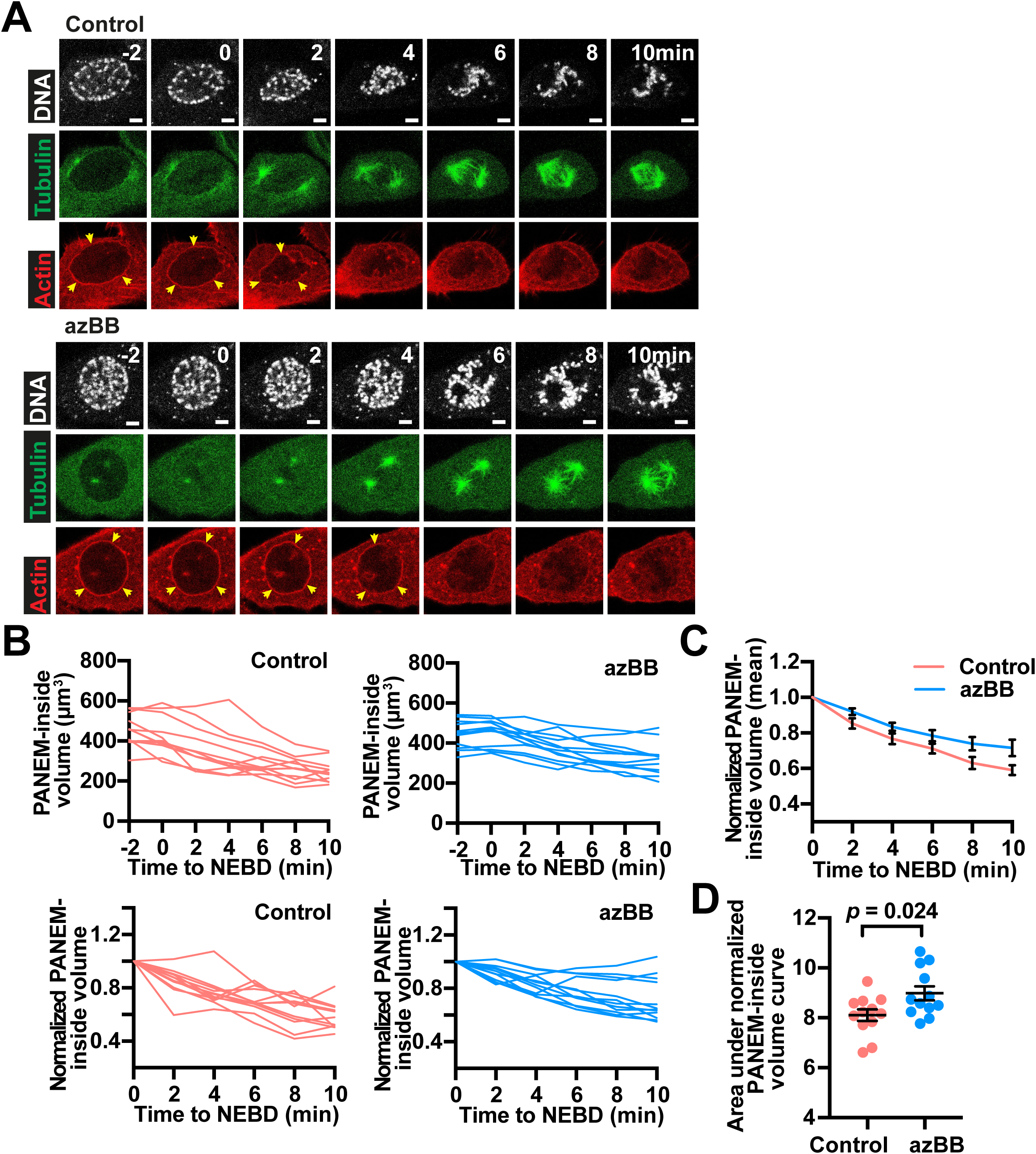
PANEM reduces its inside volume soon after NEBD. **(A)** Time-lapse images show representative cells passing through the early stages of mitosis (late prophase to prometaphase). A stable cell line expressing mCherry-LifeAct and GFP-⍰Tubulin, with chromosomes visualized by SiRDNA, was treated (or not) with azBB and imaged every 2 minutes. Times are relative to NEBD. The PANEM is indicated by yellow arrowheads. Scale bars are 5µm. **(B)** Graphs show PANEM-inside volume before and after NEBD (0 min) for cells treated with or without azBB (upper panels). Measurements were made from cells exemplified in A. In the lower panels the same data is normalized to the volume at 0 minutes (immediately after NEBD). Each red or blue line represents the measurements from an individual cell. **(C)** The mean of normalized values obtained in B for cells treated with or without azBB are presented here with SEM shown for each timepoint for each condition. **(D)** The areas under the curves for normalized PANEM-inside volume measurements in individual cells from data presented in B are shown here. The bars represent the mean and SEM. The *p* value obtained using t-test. Number of cells analyzed was 12 for both control and azBB treated cells.

**Figure 1 – figure supplement 6:**
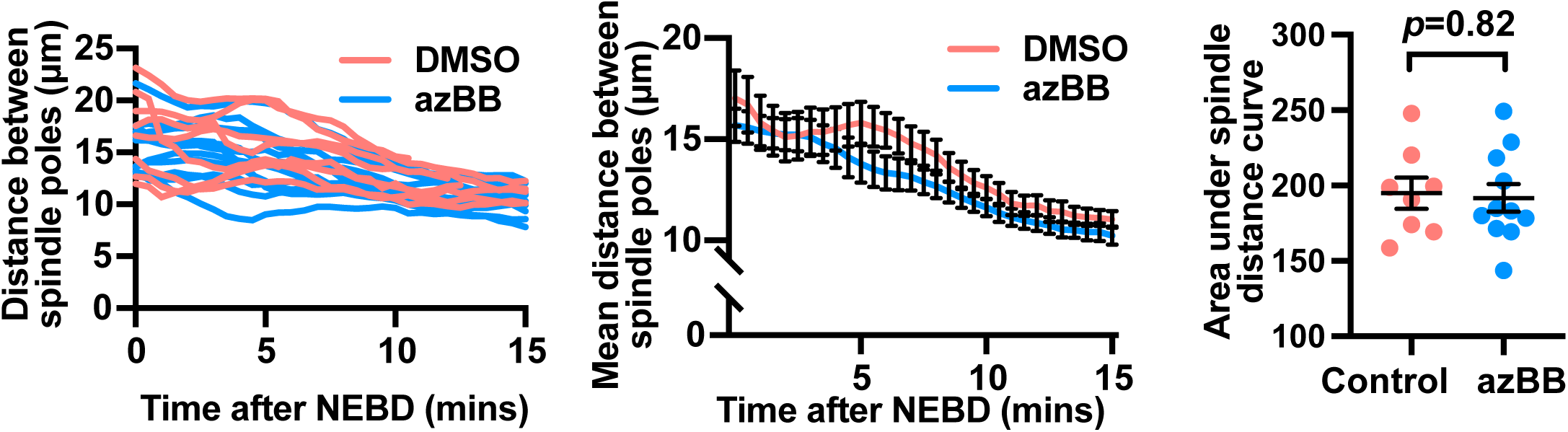
Distance between spindle poles is not affected by azBB treatment soon after NEBD. On the left-hand side, a graph shows changes in distance between spindle poles of individual cells treated with (blue lines) or without azBB (red lines) during prometaphase. In the middle, the mean values are shown with bars representing the SEM. In these graphs, times are relative to NEBD. Measurements were made from cells imaged as in Figure 2B and D. We did not observe a significant difference in the spindle-pole distance between control and azBB-treated cells at any individual time points (the smallest p-value was 0.094 at 6 min). On the right-hand side, the areas under the curves for individual cells (left-hand side graph) are shown. The bars represent the mean and SEM. The *p* value obtained using t-test. Number of cells analyzed was 8 and 11 for control and azBB treated cells, respectively.

**Figure 2 – figure supplement 1:**
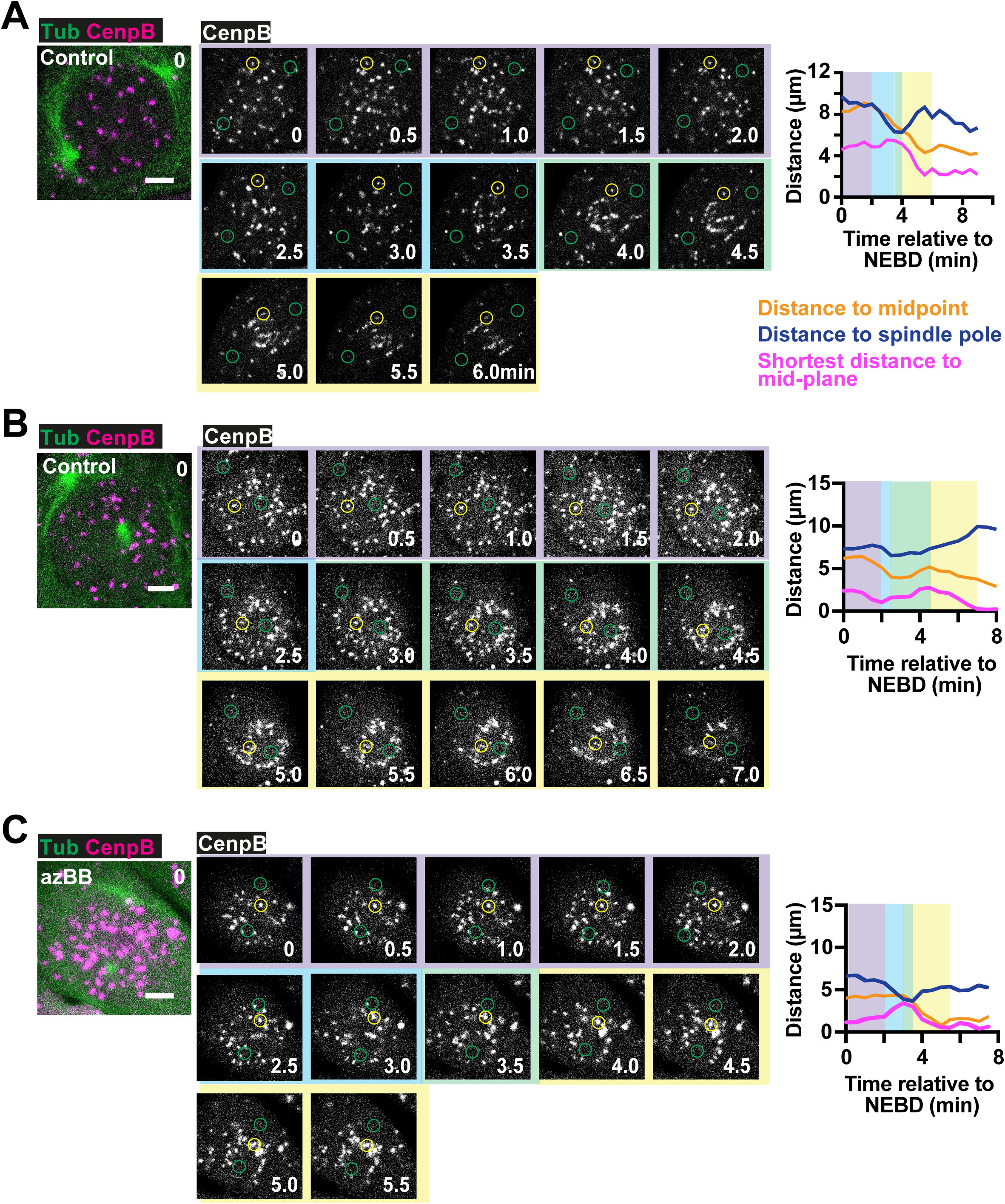
Examples of kinetochore tracking during the early stages of mitosis (prometaphase). (A-C) Time-lapse images show representative cells passing through the early stages of mitosis (prometaphase). A stable cell line expressing CENPB-mCherry and GFP-⍰Tubulin was treated with or without azBB and imaged every 30 seconds. The merged images on the left-hand side show the cells of interest at NEBD (0min). In each case the central images show the changing position of the selected kinetochores, which were tracked through time. Here the xy positions of the spindle poles are indicated by green circles while the tracked kinetochore is indicated in each frame by a yellow circle. For each time point, a single z-stack (or two z-stacks with maximum projection) containing the kinetochore of interest is shown. Scale bar is 5µm. The graph on the right-hand side indicates the distance of the indicated kinetochore to the nearest spindle pole (dark blue line), the spindle midpoint (orange line) and the spindle mid-plane (magenta line). The colored boxes on the graph, and surrounding the central images, represent the different phases of motion (explained in Figure 2C). For part A, selected time points of this time-lapse sequence for this cell, with maximum projection of up to 10 z stacks for each time point, are shown in Figure 2B. In part A, the same graph shown in Figure 2B is reproduced here. In part C, in order to show the region of the cell containing both the kinetochore of interest and the spindle poles, the merged image on the left-hand side is a maximum projection of 6 z slices.

**Figure 3 – source data 1: Raw coordinate data for peripheral kinetochores tracked in control cells.** Contains xyz coordinate data relative to time for spindle poles and peripheral kinetochores of control cells used for the analyses. The file is organised with the data from each cell grouped together. It also includes a single tab showing the timing of the different phases (described in Figure 2) for each kinetochore mentioned.

**Figure 3 – source data 2: Raw coordinate data for peripheral kinetochores tracked in azBB-treated cells.** Contains xyz coordinate data relative to time for spindle poles and peripheral kinetochores of azBB-treated cells used for the analyses. The file is organised with the data from each cell grouped together. It also includes a single tab showing the timing of the different phases (described in Figure 2) for each kinetochore mentioned.

**Figure 3 – source data 3: Raw coordinate data for central kinetochores tracked in control cells.** Contains xyz coordinate data relative to time for spindle poles and central kinetochores of control cells used for the analyses. The file is organised with the data from each cell grouped together. It also includes a single tab showing the timing of the different phases (described in Figure 2) for each kinetochore mentioned.

**Figure 3 – source data 4: Raw coordinate data for central kinetochores tracked in azBB-treated cells.** Contains xyz coordinate data relative to time for spindle poles and central kinetochores of azBB-treated cells used for the analyses. The file is organised with the data from each cell grouped together. It also includes a single tab showing the timing of the different phases (described in Figure 2) for each kinetochore mentioned.

**Figure 4 – figure supplement 1:**
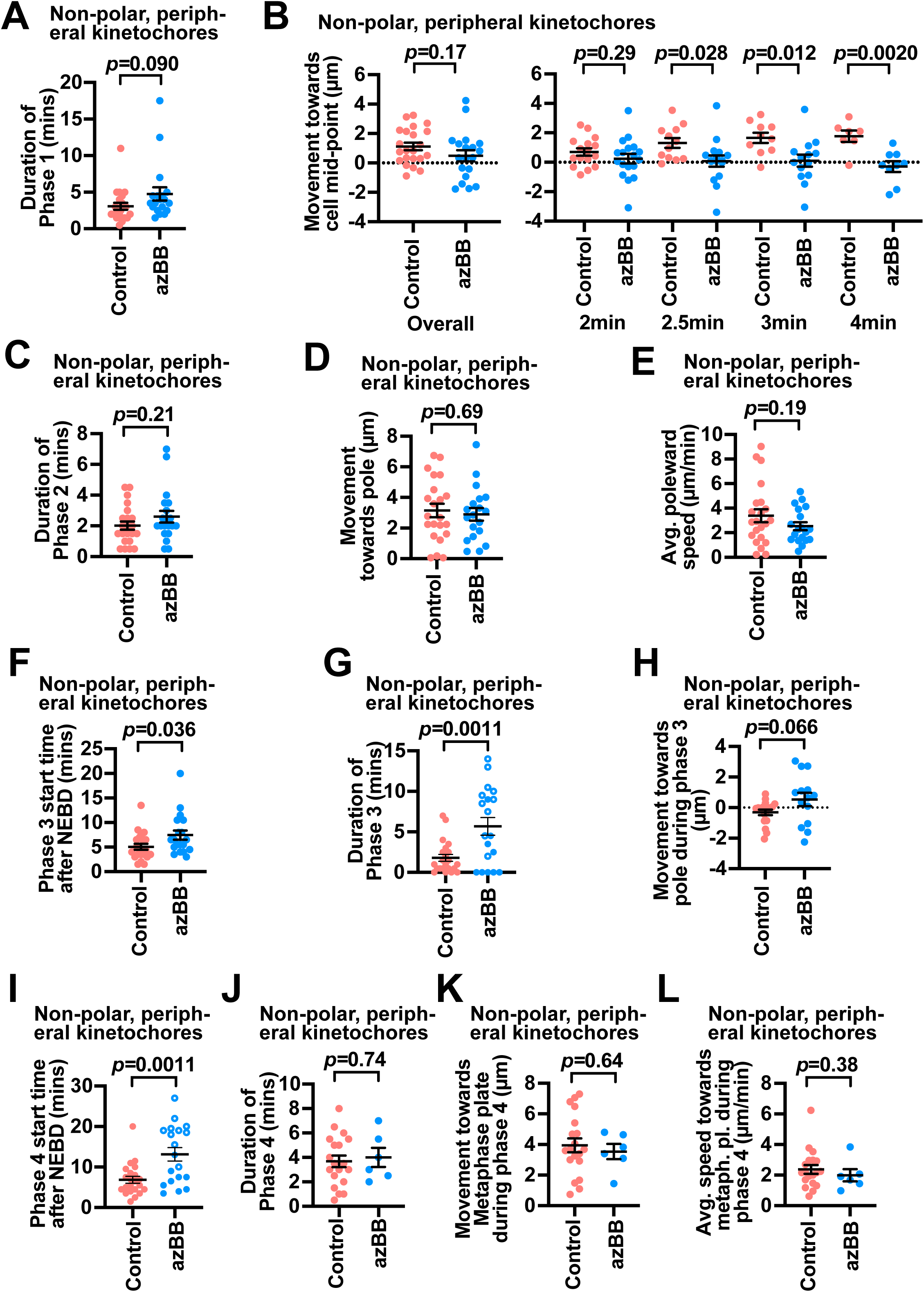
Comparison of non-polar peripheral kinetochore dynamics during each phase between control and azBB treated cells. (A–E) Graphs show the dynamics of non-polar peripheral kinetochores during Phase 1 and 2. The results were analyzed and presented as in Figure 3C–G, using the same data set, except that polar peripheral kinetochores were excluded. 22 and 19 kinetochores were analyzed from control and azBB treated cells, respectively. The *p* values were obtained by t-test. A Mann-Whitney test was also performed on data for part A giving a *p* value of 0.044. **(F–L)** Graphs show the dynamics of non-polar peripheral kinetochores during Phase 3 and 4. The results were analyzed and presented as in Figure 4B–H, using the same data set, except that polar peripheral kinetochores were excluded. The *p* values were obtained by t-test. 22 and 19 kinetochores (from 9 and 9 cells) (F–I) and 19 and 6 kinetochores (from the corresponding cells) (J–L) were analyzed from control and azBB treated cells, respectively.

**Figure 5 – figure supplement 1:**
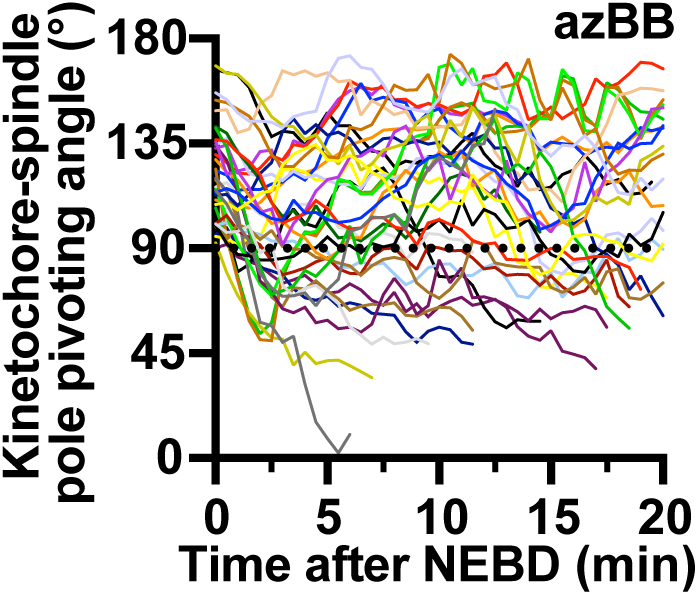
Further examples of kinetochore-spindle pole pivoting angles in azBB-treated cells. Plot shows changes in pivot angles (defined as in Figure 2A) of polar kinetochores (at NEBD) over time after NEBD (time 0), from 3 different cells treated with azBB. Individual colored lines indicate individual kinetochores. The dotted line indicates the angle at which a polar kinetochore (>90°) passed into the non-polar (central) region (<90°). The number of polar kinetochores analyzed was 36. This graph contains the data from Figure 5D.

**Figure 5 – figure supplement 2:**
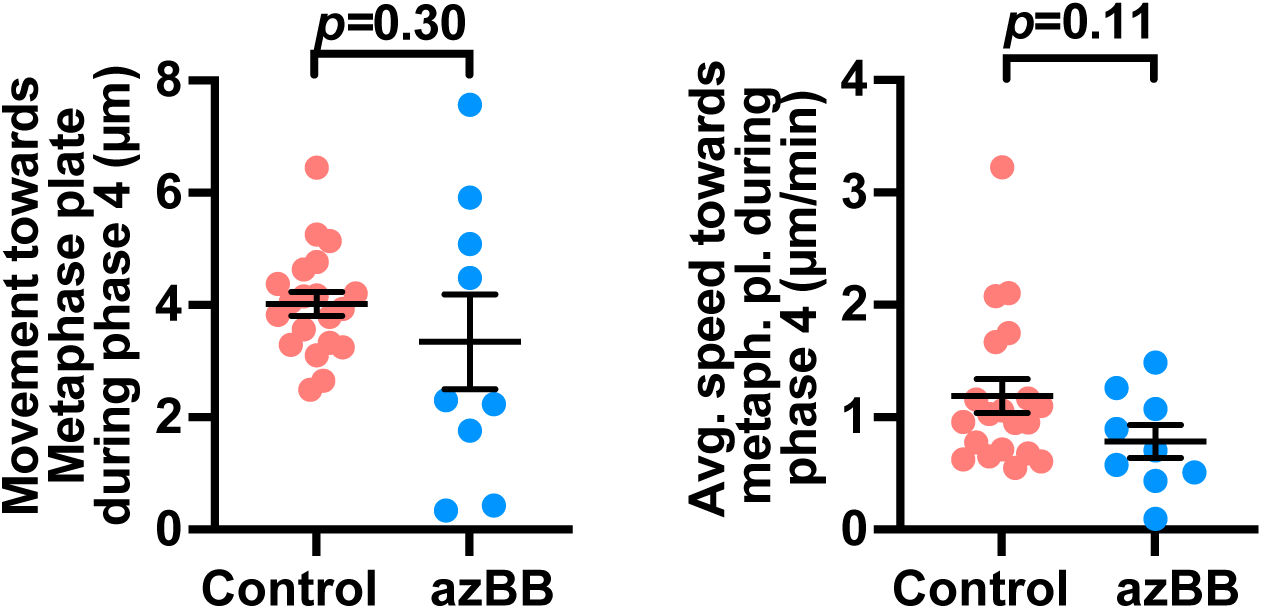
Dynamics of congression itself are not significantly affected by inhibition of PANEM contraction. Graphs show the movement of polar kinetochores towards the spindle mid-plane during congression (left-hand side) or the average speed of motion towards the spindle mid-plane during congression (right-hand side) for individual polar kinetochores from control (red) or from azBB-treated (blue) cells. The *p* values were obtained by t-test. Number of polar kinetochores was 20 and 9 from 5 control and 3 azBB-treated cells, respectively. For azBB-treated cells, graph includes only the polar kinetochores that subsequently showed congression. The analyzed kinetochores are the same as those in Figure 5F.

**Figure 5 – figure supplement 3:**
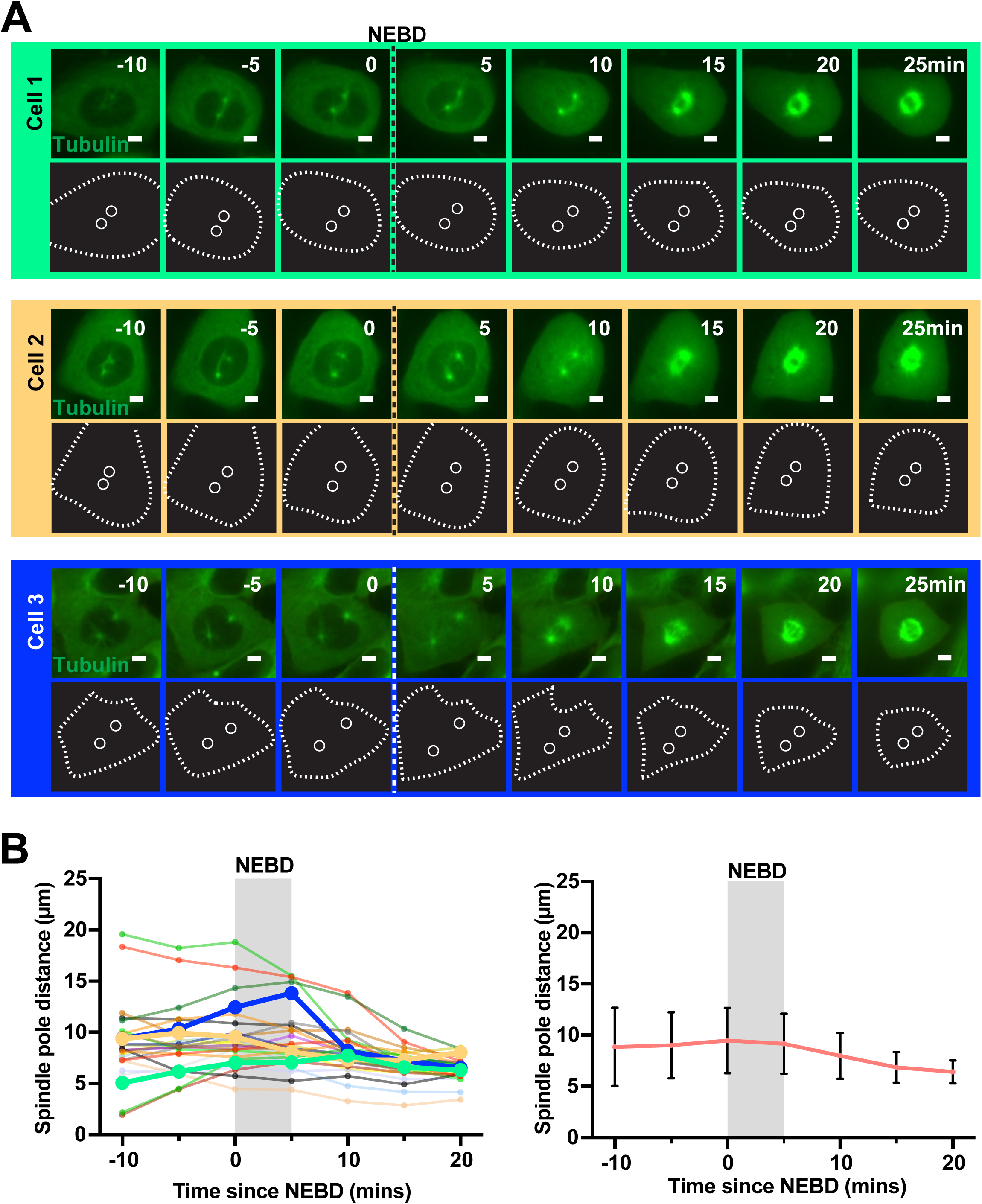
The distance between the spindle poles of asynchronous U2OS cells rarely grow following NEBD. **(A)** Time-lapse images show three representative cells passing through the early stages of mitosis (prophase and prometaphase) each one framed in a different color. A stable cell line expressing GFP-⍰Tubulin was imaged every 5 minutes. The timing of nuclear envelope breakdown (NEBD) is indicated by the dotted line. The cartoons below the images show the determined positions of the spindle poles (white circles) and the outline of the cells (white dashed lines). Scale bar is 5µm. **(B)** Graph on the left-hand side shows the distances between spindle poles over time with each line representing an individual cell. The distances of the spindle poles of the cells shown in A are the same color as the frames in A and represented with heavy lines and closed circles. On the right-hand side the mean distance for all the data is shown with bars representing the SD. Number of cells measured is 26 cells.

**Figure 5 – source data 1: Raw coordinate data for polar kinetochores tracked in control cells.** Contains xyz coordinate data relative to time for spindle poles and polar kinetochores of control cells used for the analyses. The file is organised with the data from each cell grouped together. It also includes a single tab showing the timing of the start and end of congression for each kinetochore mentioned.

**Figure 5 – source data 2: Raw coordinate data for polar kinetochores tracked in azBB-treated cells.** Contains xyz coordinate data relative to time for spindle poles and polar kinetochores of azBB-treated cells used for the analyses. The file is organised with the data from each cell grouped together. It also includes a single tab showing the timing of the start and end of congression for each kinetochore mentioned.

**Figure 6 – figure supplement 1:**
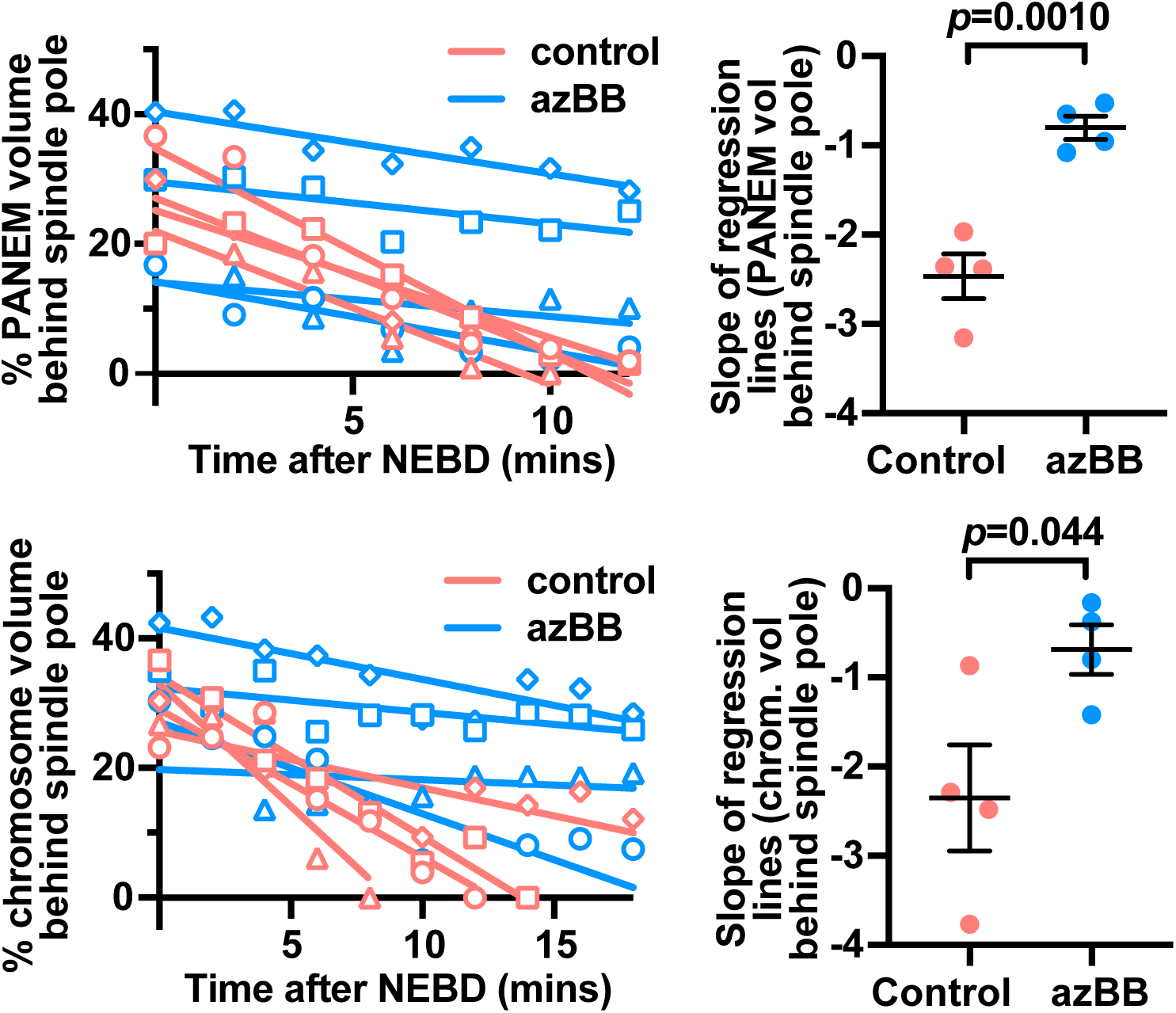
PANEM contraction is important to reduce PANEM or chromosome volume behind the spindle poles following NEBD. The left-hand side. graphs show linear regression analysis of changes in PANEM-inside volumes (upper panel) or chromosome volumes (lower panel) over time, behind the spindle poles, for individual control (red lines and shapes) or azBB-treated (blue lines and shapes) cells. The same data as in Figure 6B and C was analyzed. The right-hand side shows the slopes of the lines plotted for control (red) or azBB treated (blue) cells. The bars represent the mean and SEM. The *p* value was obtained by t-test.

**Figure 7 – supplemental figure 1:**
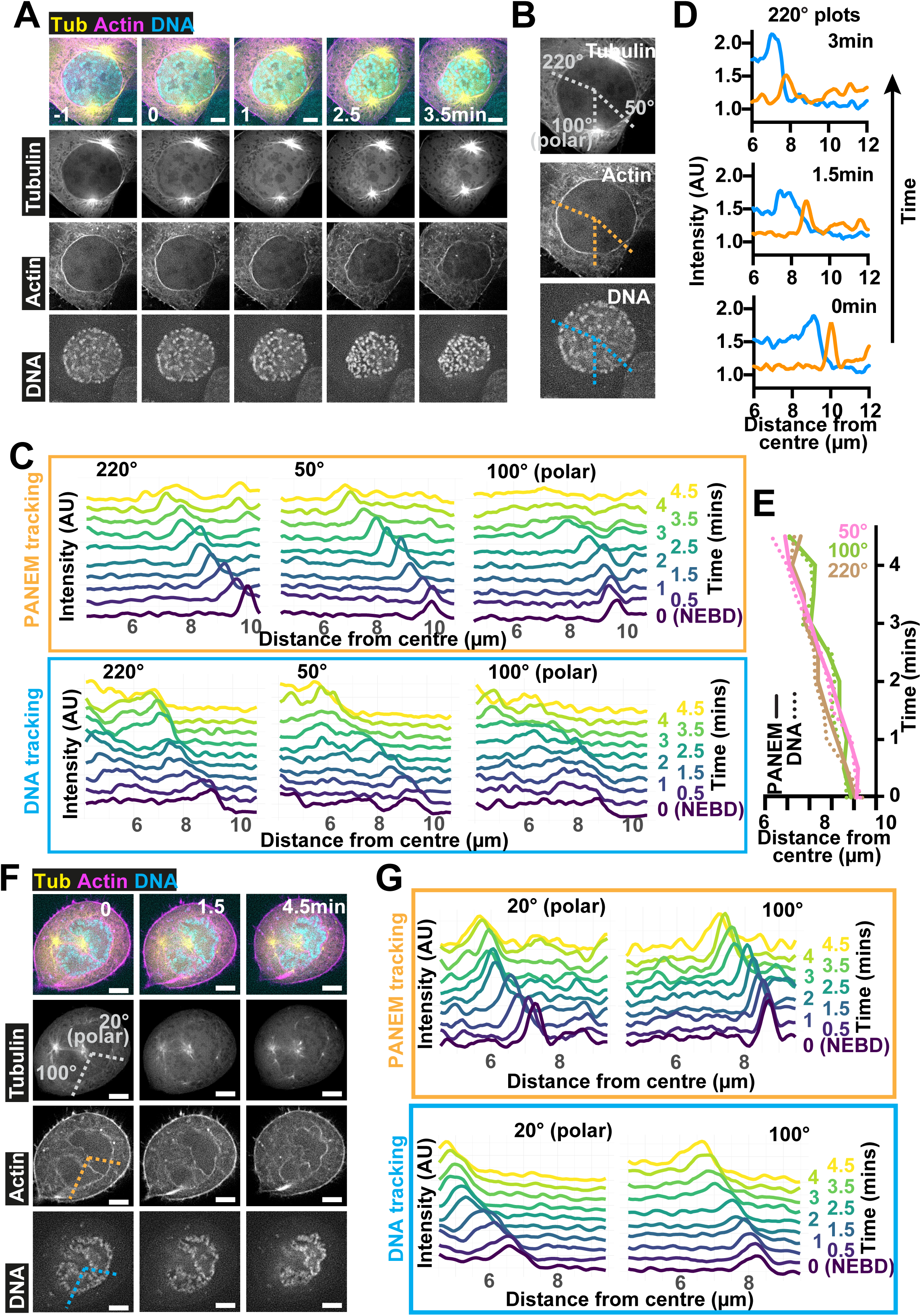
More examples showing that the contractile PANEM directly pushes both polar and non-polar peripheral chromosomes inward during early stages of mitosis. **(A)** Time-lapse images show another cell passing through the early stages of mitosis. Images were acquired and presented as in Figure 7A. NEBD is indicated at time 0. Scale bars 5µm. **(B)** Images 1 minute before NEBD from the time-lapse sequence in A to indicate the positions of line profiles. Dotted lines, plotted from the cell center (for determination see Materials and Methods), indicate the line profiles that pass the non-polar region of the cell (50° and 220°) and the polar region (100°) of the cell. In this cell, both spindle poles were close to the PANEM so there is no polar volume. **(C)** Graphs show line profiles, offset in the y-axis according to time, for PANEM (upper panel; orange frame) or chromosomes (lower panel; blue frame) for the lines indicated in B. As time progresses, peaks move to the left, which indicates movement closer towards the chromosome mass center. **(D)** Graphs show a time sequence of intensities calculated along the line profiles that pass a non-polar region of the cell shown in A. Time progresses upwards and the color of the lines indicates the intensities seen for Actin (PANEM; orange) or chromosomes (blue). **(E)** Graph shows the progression of the relationship between the PANEM peak and the chromosome front, over time, through the line profiles indicated in B. **(F)** Time-lapse images show a representative cell passing through the early stages of mitosis. Images were acquired and presented as in Figure 7A. In this cell one of the poles is close to the PANEM so there is only one pole displaying polar volume. NEBD is indicated at time 0. Scale bars 5µm. **(G)** Graphs show line profiles, offset in the y-axis according to time, for PANEM (upper panel; orange frame) or chromosomes (lower panel; blue frame) for the lines indicated in F. As time progresses, peaks move to the left, which indicates movement closer towards the chromosome mass center.

**Figure 8 – figure supplement 1:**
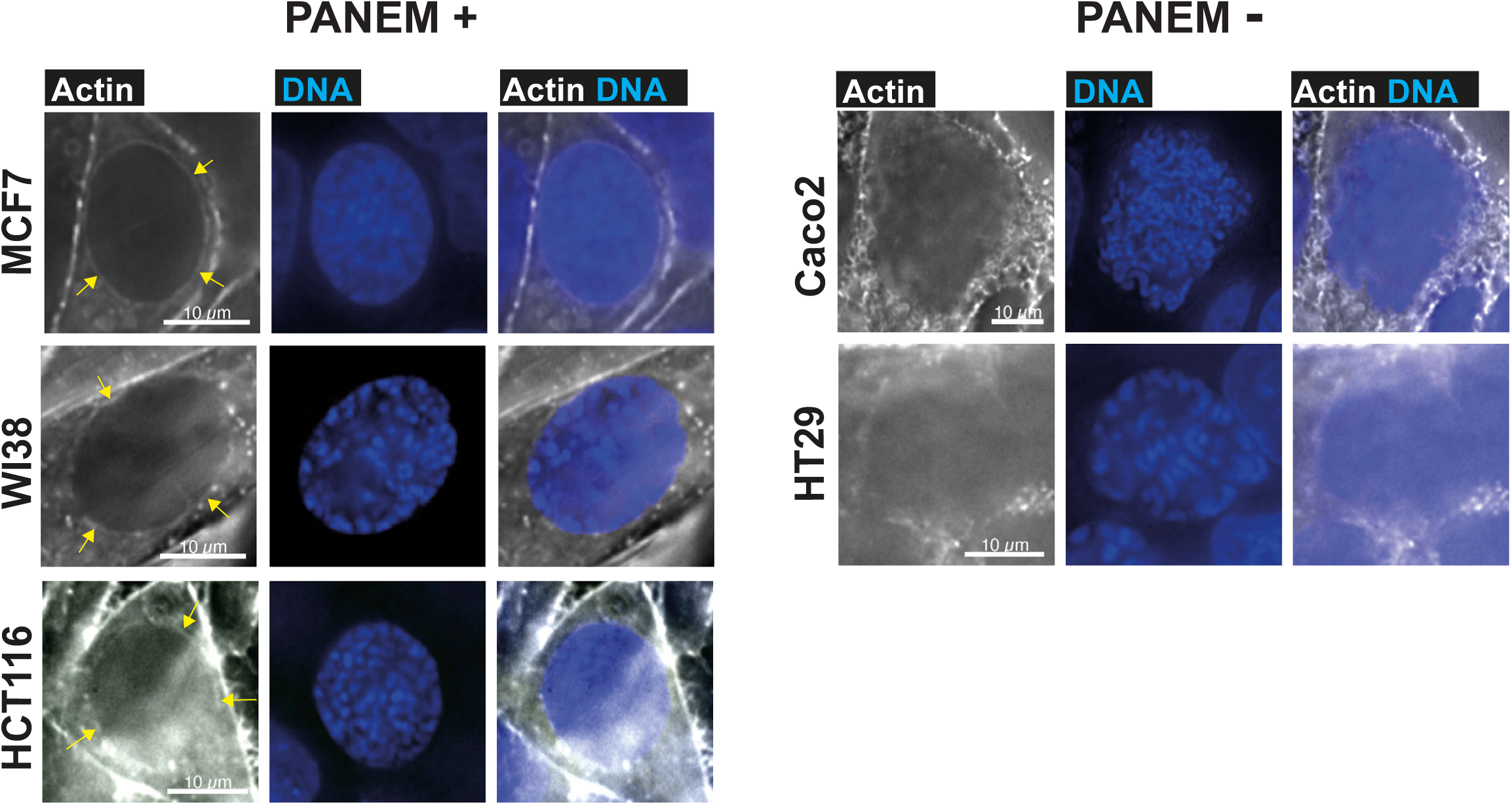
Examination of several human cancer cell lines for the presence or absence of PANEM during prophase. Images show representative prophase cells of five cancer cell lines. The cancer cell lines were fixed, and Actin (left) and DNA (middle) were visualized using phalloidin and Hoechst, respectively. Actin/DNA merged images are also shown (right). Prophase cells were identified by visualising early stages of chromosome condensation. PANEM was identified in the left-hand side panel of cells (indicated by yellow arrowheads), but not identified in the right-hand panel of cells.

**Figure 8 – figure supplement 2:**
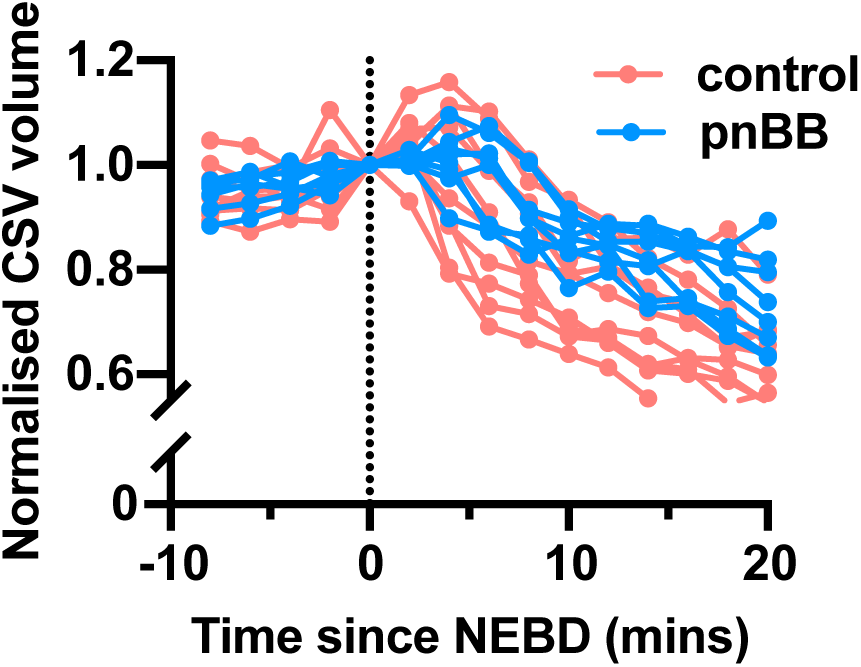
PANEM contraction is required for reducing chromosome scattering volume in the non-cancerous RPE1 cell line. Graph shows chromosome scattering volume (CSV) before and after NEBD (0 min) for RPE1 cells treated with (blue) or without (red) pnBB. Here the data from each cell is normalized to the volume at 0 minutes (immediately after NEBD). Each red or blue line represents the measurements from an individual cell. The mean for each condition is shown in Figure 8C (left hand side). The number of cells for each group was 10 and 8 for control and pnBB, respectively.

**Figure 9 – figure supplement 1:**
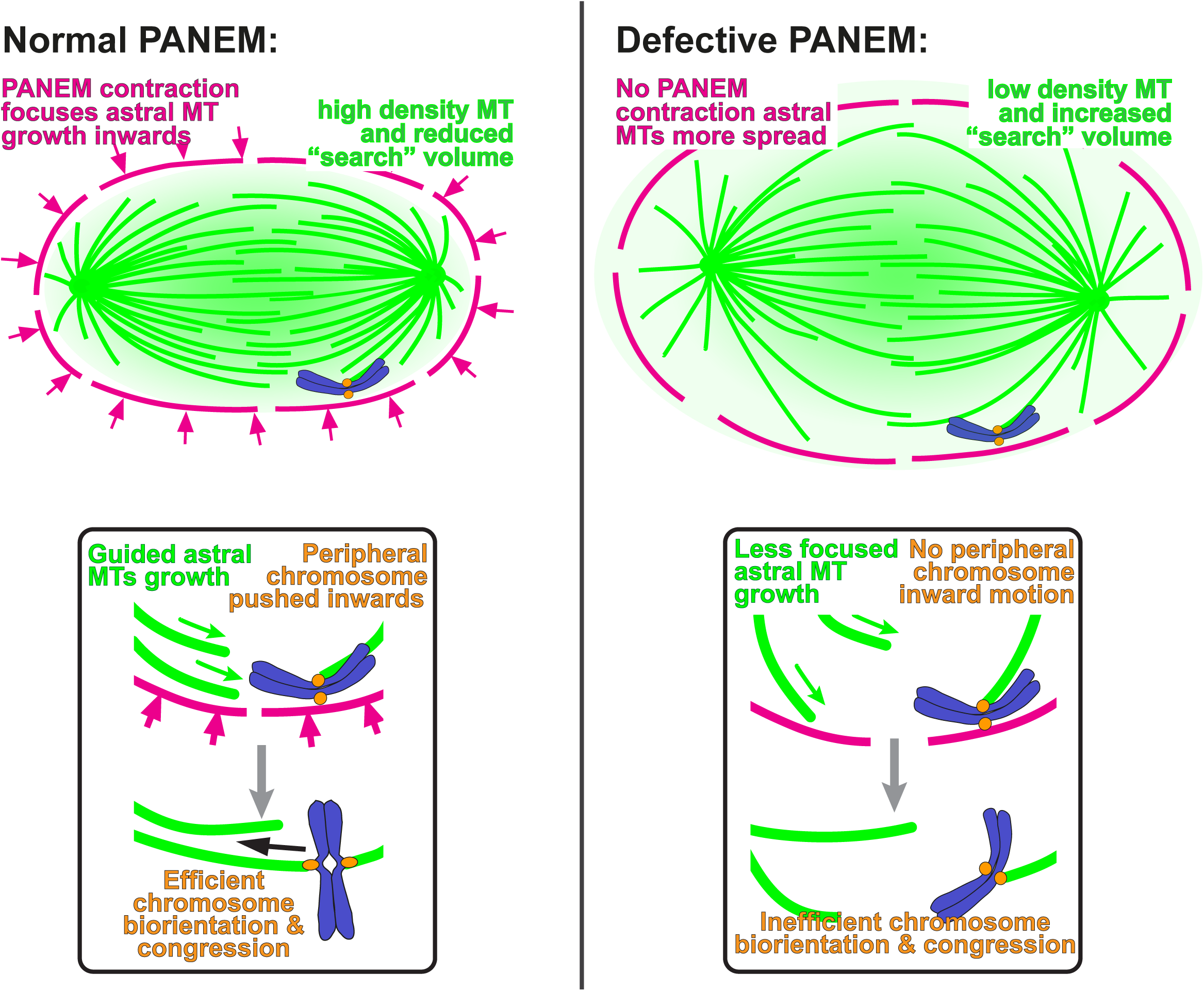
Model showing the effect of PANEM on peripheral/polar chromosomes in the situation where biorientation precedes congression. Left-hand side shows a model of how the PANEM contraction helps to prevent chromosome misalignment by pushing a peripheral chromosome inward to a region of the cell with increased density of astral microtubules emanating from the opposite cell pole. The box below indicates how directed growth of astral microtubules promote efficient capture and biorientation before congression to the cell mid plane. With defective PANEM contraction (right-hand side), peripheral chromosomes may remain in regions with lower density of astral microtubules form the opposite cell pole. The box below indicates how less focused growth leads to less efficient capture and biorientation.

## Notes

### Summary of Updates

We have revised the manuscript in response to the additional comments from one of the reviewers. We intend this to be the final version (version of record) for publication in eLife.

## References

1. Kapoor, T.M. (2017). Metaphase Spindle Assembly. Biology 6. 39–74. 10.3390/biology6010008.

2. Santaguida, S., and Musacchio, A. (2009). The life and miracles of kinetochores. EMBO J 28, 2511–2531. 10.1038/emboj.2009.173

3. Hinshaw, S.M., and Harrison, S.C. (2018). Kinetochore Function from the Bottom Up. Trends Cell Biol 28, 22–33. 10.1016/j.tcb.2017.09.002.

4. Musacchio, A., and Desai, A. (2017). A Molecular View of Kinetochore Assembly and Function. Biology (Basel) 6. 75–121 10.3390/biology6010005.

5. Magidson, V., O’Connell, C.B., Loncarek, J., Paul, R., Mogilner, A., and Khodjakov, A. (2011). The spatial arrangement of chromosomes during prometaphase facilitates spindle assembly. Cell 146, 555–567. 10.1016/j.cell.2011.07.012.

6. Tanaka, T.U., and Zhang, T. (2022). SWAP, SWITCH, and STABILIZE: Mechanisms of Kinetochore-Microtubule Error Correction. Cells 11. 1462. 10.3390/cells11091462.

7. Rieder, C.L., and Alexander, S.P. (1990). Kinetochores are transported poleward along a single astral microtubule during chromosome attachment to the spindle in newt lung cells. J Cell Biol 110, 81–95.

8. Tanaka, K., Kitamura, E., Kitamura, Y., and Tanaka, T.U. (2007). Molecular mechanisms of microtubule-dependent kinetochore transport toward spindle poles. J Cell Biol 178, 269–281. 10.1083/jcb.200702141.

9. Tanaka, K., Mukae, N., Dewar, H., van Breugel, M., James, E.K., Prescott, A.R., Antony, C., and Tanaka, T.U. (2005). Molecular mechanisms of kinetochore capture by spindle microtubules. Nature 434, 987–994.

10. Shrestha, R.L., and Draviam, V.M. (2013). Lateral to end-on conversion of chromosome-microtubule attachment requires kinesins CENP-E and MCAK. Curr Biol. 23, 1514–1526. doi: 1510.1016/j.cub.2013.1506.1040. Epub 2013 Jul 1525.

11. Itoh, G., Ikeda, M., Iemura, K., Amin, M.A., Kuriyama, S., Tanaka, M., Mizuno, N., Osakada, H., Haraguchi, T., and Tanaka, K. (2018). Lateral attachment of kinetochores to microtubules is enriched in prometaphase rosette and facilitates chromosome alignment and bi-orientation establishment. Scientific reports 8, 3888. 10.1038/s41598-018-22164-5.

12. Kapoor, T.M., Lampson, M.A., Hergert, P., Cameron, L., Cimini, D., Salmon, E.D., McEwen, B.F., and Khodjakov, A. (2006). Chromosomes can congress to the metaphase plate before biorientation. Science 311, 388–391.

13. Maiato, H., Gomes, A.M., Sousa, F., and Barisic, M. (2017). Mechanisms of Chromosome Congression during Mitosis. Biology 6. 10.3390/biology6010013.

14. Godek, K.M., Kabeche, L., and Compton, D.A. (2015). Regulation of kinetochore-microtubule attachments through homeostatic control during mitosis. Nat Rev Mol Cell Biol 16, 57–64. 10.1038/nrm3916.

15. Armond, J.W., Harry, E.F., McAinsh, A.D., and Burroughs, N.J. (2015). Inferring the Forces Controlling Metaphase Kinetochore Oscillations by Reverse Engineering System Dynamics. PLoS Comput Biol 11, e1004607. 10.1371/journal.pcbi.1004607.

16. Jaqaman, K., King, E.M., Amaro, A.C., Winter, J.R., Dorn, J.F., Elliott, H.L., McHedlishvili, N., McClelland, S.E., Porter, I.M., Posch, M., et al. (2010). Kinetochore alignment within the metaphase plate is regulated by centromere stiffness and microtubule depolymerases. The Journal of cell biology 188, 665–679. 10.1083/jcb.200909005.

17. Skibbens, R.V., Skeen, V.P., and Salmon, E.D. (1993). Directional instability of kinetochore motility during chromosome congression and segregation in mitotic newt lung cells: a push-pull mechanism. J Cell Biol 122, 859–875.

18. Vukusic, K., and Tolic, I.M. (2022). Polar Chromosomes-Challenges of a Risky Path. Cells 11. 10.3390/cells11091531.

19. Klaasen, S.J., Truong, M.A., van Jaarsveld, R.H., Koprivec, I., Stimac, V., de Vries, S.G., Risteski, P., Kodba, S., Vukusic, K., de Luca, K.L., et al. (2022). Nuclear chromosome locations dictate segregation error frequencies. Nature 607, 604–609. 10.1038/s41586-022-04938-0.

20. Booth, A.J.R., Yue, Z., Eykelenboom, J.K., Stiff, T., Luxton, G.W.G., Hochegger, H., and Tanaka, T.U. (2019). Contractile acto-myosin network on nuclear envelope remnants positions human chromosomes for mitosis. eLife 8. 10.7554/eLife.46902.

21. Stiff, T., Echegaray-Iturra, F.R., Pink, H.J., Herbert, A., Reyes-Aldasoro, C.C., and Hochegger, H. (2020). Prophase-Specific Perinuclear Actin Coordinates Centrosome Separation and Positioning to Ensure Accurate Chromosome Segregation. Cell reports 31, 107681. 10.1016/j.celrep.2020.107681.

22. Barisic, M., Aguiar, P., Geley, S., and Maiato, H. (2014). Kinetochore motors drive congression of peripheral polar chromosomes by overcoming random arm-ejection forces. Nature cell biology 16, 1249–1256. 10.1038/ncb3060.

23. Koprivec, I., Stimac, V., Dura, M., Vukusic, K., Mikec, P., and Tolic, I.M. (2026). Polar chromosomes are rescued from missegregation by spindle elongation-driven microtubule pivoting. Nature communications 17. 10.1038/s41467-026-69830-1.

24. Renda, F., Miles, C., Tikhonenko, I., Fisher, R., Carlini, L., Kapoor, T.M., Mogilner, A., and Khodjakov, A. (2022). Non-centrosomal microtubules at kinetochores promote rapid chromosome biorientation during mitosis in human cells. Current biology : CB 32, 1049–1063 e1044. 10.1016/j.cub.2022.01.013.

25. Vukusic, K., and Tolic, I.M. (2025). CENP-E initiates chromosome congression by opposing Aurora kinases to promote end-on attachments. Nature communications 16, 8537. 10.1038/s41467-025-64148-w.

26. Kepiro, M., Varkuti, B.H., Bodor, A., Hegyi, G., Drahos, L., Kovacs, M., and Malnasi-Csizmadia, A. (2012). Azidoblebbistatin, a photoreactive myosin inhibitor. Proceedings of the National Academy of Sciences of the United States of America 109, 9402–9407. 10.1073/pnas.1202786109.

27. Képiró, M., Várkuti, Boglárka H., Rauscher, Anna A., Kellermayer, Miklós S.Z., Varga, M., and Málnási-Csizmadia, A. (2015). Molecular Tattoo: Subcellular Confinement of Drug Effects. Chemistry & Biology 22, 548–558. 10.1016/j.chembiol.2015.03.013.

28. Taubenberger, A.V., Baum, B., and Matthews, H.K. (2020). The Mechanics of Mitotic Cell Rounding. Front Cell Dev Biol 8, 687. 10.3389/fcell.2020.00687.

29. Devillers, R., Dos Santos, A., Destombes, Q., Laplante, M., and Elowe, S. (2024). Recent insights into the causes and consequences of chromosome mis-segregation. Oncogene 43, 3139–3150. 10.1038/s41388-024-03163-5.

30. Aquino-Perez, C., Safaralizade, M., Podhajecky, R., Wang, H., Lansky, Z., Grosse, R., and Macurek, L. (2024). FAM110A promotes mitotic spindle formation by linking microtubules with actin cytoskeleton. Proceedings of the National Academy of Sciences of the United States of America 121, e2321647121. 10.1073/pnas.2321647121.

31. Kita, A.M., Swider, Z.T., Erofeev, I., Halloran, M.C., Goryachev, A.B., and Bement, W.M. (2019). Spindle-F-actin interactions in mitotic spindles in an intact vertebrate epithelium. Molecular biology of the cell 30, 1645–1654. 10.1091/mbc.E19-02-0126.

32. Plessner, M., Knerr, J., and Grosse, R. (2019). Centrosomal Actin Assembly Is Required for Proper Mitotic Spindle Formation and Chromosome Congression. iScience 15, 274–281. 10.1016/j.isci.2019.04.022.

33. Limouze, J., Straight, A.F., Mitchison, T., and Sellers, J.R. (2004). Specificity of blebbistatin, an inhibitor of myosin II. J Muscle Res Cell Motil 25, 337–341. 10.1007/s10974-004-6060-7.

34. Mayca Pozo, F., Geng, X., Miyagi, M., Amin, A.L., Huang, A.Y., and Zhang, Y. (2023). MYO10 regulates genome stability and cancer inflammation through mediating mitosis. Cell reports 42, 112531. 10.1016/j.celrep.2023.112531.

35. Woolner, S., O’Brien, L.L., Wiese, C., and Bement, W.M. (2008). Myosin-10 and actin filaments are essential for mitotic spindle function. The Journal of cell biology 182, 77–88.

36. Kaseda, K., McAinsh, A.D., and Cross, R.A. (2012). Dual pathway spindle assembly increases both the speed and the fidelity of mitosis. Biol Open 1, 12–18. 10.1242/bio.2011012.

37. Lombardi, M.L., and Lammerding, J. (2011). Keeping the LINC: the importance of nucleocytoskeletal coupling in intracellular force transmission and cellular function. Biochem Soc Trans 39, 1729–1734. 10.1042/BST20110686.

38. Firat-Karalar, E.N., and Welch, M.D. (2011). New mechanisms and functions of actin nucleation. Current opinion in cell biology 23, 4–13. 10.1016/j.ceb.2010.10.007.

39. Beaudouin, J., Gerlich, D., Daigle, N., Eils, R., and Ellenberg, J. (2002). Nuclear envelope breakdown proceeds by microtubule-induced tearing of the lamina. Cell 108, 83–96. 10.1016/s0092-8674(01)00627-4.

40. Dewey, E.B., and Johnston, C.A. (2017). Diverse mitotic functions of the cytoskeletal cross-linking protein Shortstop suggest a role in Dynein/Dynactin activity. Molecular biology of the cell 28, 2555–2568. 10.1091/mbc.E17-04-0219.

41. Lancaster, O.M., Le Berre, M., Dimitracopoulos, A., Bonazzi, D., Zlotek-Zlotkiewicz, E., Picone, R., Duke, T., Piel, M., and Baum, B. (2013). Mitotic rounding alters cell geometry to ensure efficient bipolar spindle formation. Developmental cell 25, 270–283. 10.1016/j.devcel.2013.03.014.

42. di Pietro, F., Echard, A., and Morin, X. (2016). Regulation of mitotic spindle orientation: an integrated view. EMBO Rep 17, 1106–1130. 10.15252/embr.201642292.

43. Kunda, P., and Baum, B. (2009). The actin cytoskeleton in spindle assembly and positioning. Trends Cell Biol 19, 174–179. 10.1016/j.tcb.2009.01.006.

44. Pollard, T.D., and O’Shaughnessy, B. (2019). Molecular Mechanism of Cytokinesis. Annual review of biochemistry 88, 661–689. 10.1146/annurev-biochem-062917-012530.

45. Davidson, P.M., and Cadot, B. (2021). Actin on and around the Nucleus. Trends Cell Biol 31, 211–223. 10.1016/j.tcb.2020.11.009.

46. Burdyniuk, M., Callegari, A., Mori, M., Nédélec, F., and Lénárt, P. (2018). F-Actin nucleated on chromosomes coordinates their capture by microtubules in oocyte meiosis. The Journal of cell biology 217, 2661–2674.

47. Lenart, P., Bacher, C.P., Daigle, N., Hand, A.R., Eils, R., Terasaki, M., and Ellenberg, J. (2005). A contractile nuclear actin network drives chromosome congression in oocytes. Nature 436, 812–818. 10.1038/nature03810.

48. Harasimov, K., Uraji, J., Monnich, E.U., Holubcova, Z., Elder, K., Blayney, M., and Schuh, M. (2023). Actin-driven chromosome clustering facilitates fast and complete chromosome capture in mammalian oocytes. Nature cell biology 25, 439–452. 10.1038/s41556-022-01082-9.

49. Mogessie, B., and Schuh, M. (2017). Actin protects mammalian eggs against chromosome segregation errors. Science 357. 10.1126/science.aal1647.

50. Hernandez, B., Tetlak, P., Domingo-Muelas, A., Akizawa, H., Skory, R.M., Ardestani, G., Biro, M., Liu, X., Bissiere, S., Sakkas, D., and Plachta, N. (2025). Actin organizes chromosomes and microtubules to ensure mitotic fidelity in the preimplantation embryo. Science 388, eads1234. 10.1126/science.ads1234.

51. Ertych, N., Stolz, A., Stenzinger, A., Weichert, W., Kaulfuss, S., Burfeind, P., Aigner, A., Wordeman, L., and Bastians, H. (2014). Increased microtubule assembly rates influence chromosomal instability in colorectal cancer cells. Nature cell biology 16, 779–791. 10.1038/ncb2994.

52. Rata, S., Suarez Peredo Rodriguez, M.F., Joseph, S., Peter, N., Echegaray Iturra, F., Yang, F., Madzvamuse, A., Ruppert, J.G., Samejima, K., Platani, M., et al. (2018). Two Interlinked Bistable Switches Govern Mitotic Control in Mammalian Cells. Current biology 28, 3824–3832 e3826. 10.1016/j.cub.2018.09.059.

53. Arakawa, H., Lodygin, D., and Buerstedde, J.M. (2001). Mutant loxP vectors for selectable marker recycle and conditional knock-outs. BMC Biotechnol 1, 7. 10.1186/1472-6750-1-7.

54. Schneider, C.A., Rasband, W.S., and Eliceiri, K.W. (2012). NIH Image to ImageJ: 25 years of image analysis. Nature methods 9, 671–675. 10.1038/nmeth.2089.

55. Cohen-Sharir, Y., McFarland, J.M., Abdusamad, M., Marquis, C., Bernhard, S.V., Kazachkova, M., Tang, H., Ippolito, M.R., Laue, K., Zerbib, J., et al. (2021). Aneuploidy renders cancer cells vulnerable to mitotic checkpoint inhibition. Nature 590, 486–491. 10.1038/s41586-020-03114-6.

