## Supplementary material for "Contractile perinuclear actomyosin network promotes peripheral and polar chromosome interaction with the mitotic spindle": Response to reviewers

**Responses (in blue) to reviewer’s comments (in black)** 

Reviewer #1 (Recommendations for the authors): 

Overall, I believe the manuscript is suitable for publication. I have only a few minor comments and suggestions for the authors: 

(1) Some sentences remain unclear in meaning or would benefit from grammatical revision. For example: the last sentence of the Abstract (the main point not clear); the long sentence on page 6 beginning with "Although these problems might have..."; the sentence on page 3 beginning with "For example, chromosomes...," which would benefit from explicitly stating that the observation was made in tumor or otherwise perturbed cells; and the long sentence on page 9 beginning with "To avoid this potential bias...".

We have changed the text as follows:

- Abstract – this has been reworded, and the last sentence changed to make the context clearer.
- Page 3: “For example, …” – has been reworded, including a brief statement as to how the observations were made.
- Page 6: “Although these…” – we have reworded this paragraph to make clearer what we meant.
- Page 9: “To avoid this…” – We have reworded this section to make our intentions clearer for readers.

(2) When discussing the differential effects of PANEM on the phases of chromosome movement in the Results section, the conclusions would benefit from being stated more clearly in biological terms. This could be done at the end of each subsection or in a short summary paragraph (e.g., "Summarizing the analysis of Phases 1-4..."), rather than focusing primarily on the phase-based interpretation, which is already defined by underlying biological events.

Along the lines suggested by the reviewer, we have changed the second paragraph on page 9 and added a couple of sentences to the bottom paragraph on page 10.

(3) The placement of the paragraph beginning with "So far, we randomly..." on page 10 feels somewhat out of context. It may read more naturally if moved later in the Results section, when polar chromosomes are formally introduced.

We understand the reviewer's concern, but we think this analysis is best discussed at the end of this section. To clarify our intentions, we have reworded this paragraph.

(4) I could not find a citation to Figure 8-figure supplement 2 in the text.

This has now been added to the text (see page 14).

(5) Figures: The labeling could be improved for clarity. In Figure 1E, the green and orange categories could be more explicitly defined as "Completed in 30 min" and "Not completed in 30 min." A similar clarification would help in Figure 5C (e.g., "Congressed in X min"). In Figure 7, please specify from which reference point distances were measured; based on the schematic in Figure 7B and Methods, this appears to be the nuclear center. The same clarifications should be applied consistently across the supplementary figures.

- The suggested corrections have been made to Figure 1E.
- For Figure 5C, we cannot say “congressed in X minutes” because the length of time imaged after NEBD varies from cell to cell (and therefore kinetochore to kinetochore). We have added an alternative explanation to the figure legend (page 33) to clarify this point.
- For Figure 7, the reference point was the center of the cell (defined using Imaris surface function). We have added this clarification to the figure legend for Figure 7 (page 34/35) and Figure 7 supplement 1 (page 42).

(6) Please specify in the Methods (page 22) the duration of irradiation with the 860 nm laser.

We have added the requested experimental detail to the Materials and Methods (page 23)

(7) Minor figure issues: In Figure 1-figure supplement 5, panel B (top right), PANEM is mislabeled as "PANE." Please also clarify what the green signal represents in Figure 5-figure supplement 3. In the same figure, Cell 1 is missing the frame 25 image, although the schematic is present. Additionally, are the different spindle pole sizes in Figure 2A meant to reflect a functional difference, or are they just for 3D effect?

- The minor issues in Figure 1-figure supplement 5 and Figure 5-figure supplement 3 have been corrected.
- Different pole sizes in Figure 2A are just for a 3D effect.

(8) Finally, it would be valuable for the authors to discuss the variability of PANEM across single cells within the same cell line, both in terms of presence and intensity. As no quantification is currently provided, and Figure 8-figure supplement 1 shows only single-cell examples (understandably, given experimental time constraints), a brief discussion addressing the extent of cell-to-cell variability would strengthen the manuscript.

Regarding the presence of PANEM, we observed PANEM in all prophase cells (29 out of 29) in the experiment for Figure 7. We have added this information to the first paragraph on page 13. Regarding the variation of PANEM intensity, we analyzed it in individual fixed cells in our previous study (Figure 2C in Booth et al 2019). Moreover, in the current study, we analysed the variation of the PANEM inside volume among individual live cells (Figure 1- figure supplement 5).
